# Specialised RNA decay fine-tunes monogenic antigen expression in African trypanosomes

**DOI:** 10.1101/2025.05.24.654301

**Authors:** Lianne I. M. Lansink, Leon Walther, Sophie Longmore, Madeleine Jones, Adam Dowle, Joana R. C. Faria

## Abstract

Antigenic variation is a sophisticated immune evasion strategy employed by many pathogens. *Trypanosoma brucei* expresses a single Variant-Surface-Glycoprotein (VSG) from a large genetic repertoire, which they periodically switch throughout an infection. Co-transcribed with the active-*VSG* within a specialised nuclear body are expression-site-associated-genes (*ESAGs*), involved in important host-parasite interactions, including protecting the parasite from human serum lytic effects, modulating the host’s innate immune response and uptake of essential nutrients. Despite expression within the same polycistron, there is a significant differential expression between *ESAGs* and *VSGs* (>140-fold), however, the regulatory mechanism has remained elusive for decades. Here, using a combination of genetic tools, super resolution microscopy, proteomics and transcriptomics analyses, we identified three novel proteins, which are recruited in a hierarchical manner, forming discreet sub-nuclear condensates that are developmentally regulated and negatively regulate *ESAG* transcripts. Among them, Expression-Site-Body-specific-protein-2 (ESB2) contains a nuclease domain that shares structural similarity to the endonuclease domain found in SMG6, a critical component of nonsense mediated decay in mammals. Mutation of key residues required for the nuclease activity impaired ESB2 localisation and function. Overall, our findings reveal a novel mechanism of post-transcriptional regulation and shed light on how specialised RNA decay can regulate expression of specific genes.

## Introduction

The molecular mechanisms regulating expression of a single gene out of a large gene family remain major outstanding mysteries in eukaryotic biology. One of the most fascinating examples of allelic exclusion underpins the ‘arms race’ between many pathogens and their hosts through monogenic expression of surface exposed antigens and immunoglobulins, respectively [1, 2]. Among such pathogens are unicellular parasitic protozoa that cause malaria and sleeping sickness [3, 4].

African trypanosomes are transmitted by tsetse flies, in the mammalian host, they replicate freely in the bloodstream and several extra-vascular spaces, fully exposed to the host immune system [5]. They rely on antigenic variation, a highly sophisticated virulence mechanism, for successful immune evasion. To this end, they express a single variant surface glycoprotein (VSG), which they periodically switch in a stochastic manner. The active-VSG is GPI-anchored to the membrane, forming a dense coat on their surface [2, 3, 6, 7]. Notably, *VSG* expression is a remarkable example of ‘extreme biology’ whereby a single variant is expressed from the largest gene family in nature (>2,600 genes and pseudogenes), at very high levels, with the *VSG* mRNA and protein being the most abundant in the cell (5–10% of the total in each case) [8].

Despite the vast genetic repertoire, *VSG* genes are exclusively expressed from one of ∼15 sub-telomeric, polycistronic transcription-units, designated *VSG* expression-sites (*VSG*-ESs) [9]. RNA polymerase-I (Pol-I) transcribes the single active-*VSG*-ES within a specialised nuclear body, the expression-site body (ESB), while other *VSG*-ESs, despite sharing a common set of DNA elements, are transcriptionally repressed [10]. Each *VSG*-ES also contains several expression-site-associated-genes (*ESAGs*). Some are surface exposed and involved in host-parasite interactions, including protecting the parasite from human serum lytic effects, modulating the host’s innate immune response, uptake of essential nutrients and contribution to antigenic variation [11–19].

Numerous telomere-associated factors, epigenetic regulators and chromatin remodellers have been implicated in *VSG* gene silencing [20–29]. The ESB, however, has remained largely mysterious for over 20 years. Recently, through a combination of affinity purification-based approaches as well as genetic and image-based screens, significant advances have been made. Yet only two ESB components have been identified to date: ESB-specific protein 1 (ESB1) and VSG-Exclusion protein 2 (VEX2) [30, 31]. Both form discrete protein condensates in the mammalian-stage of the parasite but display distinct roles: ESB1 is a transcriptional activator whereas VEX2 prevents activation of silent ES-associated *VSGs*, rendering them positive and negative regulators of *VSG* expression, respectively [30, 31].

Notably, besides its role as an exclusion factor, VEX2 has also been shown to play a pivotal role in mediating interchromosomal interactions associated with increased *VSG* mRNA processing [32, 33]. Specifically, VEX2 associates with the chromatin at the single active *VSG*-ES and forms an allele-selective interchromosomal bridge, via VEX1 [34], to one of the two *SL*-arrays [32, 33]. These arrays encode for the *SL*-RNA required for *trans*-splicing, a process that adds a common *spliced leader* sequence to each pre-mRNA and is coupled to polyadenylation in kinetoplastid parasites [35].

Whilst Pol-I transcription hugely contributes to such high levels of *VSG* expression, several post-transcriptional mechanisms also play a critical role. On the one hand, transcription and splicing are spatially integrated in the 3D nuclear space to maximise *VSG* mRNA processing and maturation. Indeed, the ESB was not only shown to be in close spatial proximity to the *SL*-array-associated body (SLAB), but also to two other nuclear bodies involved in RNA processing and maturation - a NUFIP-body and a ‘Cajal-like’ body [32, 36]. In simple terms, a conglomerate of nuclear bodies, a ‘nuclear factory’, sustains high levels of *VSG* expression. On the other hand, both the VSG mRNA and protein are remarkably stable [37–39]. The *VSG* mRNA contains a highly conserved ’16-mer’ sequence in its 3’UTR, which binds CFB2, a cyclin-like F-box protein, and promotes N6-methyladenosine (m6A) modification of the poly(A) tail. Both m6A and CFB2 greatly contribute to the stability of the *VSG* mRNA [37, 38].

Intriguingly, it has long been known that *ESAGs* upstream of *VSGs*, despite co-transcription in the same polycistron, yield far fewer transcripts (>140-fold less), reflecting the need for much higher levels of *VSG* expression and implicating a layer of post-transcriptional regulation within the ESB that has remained elusive for decades. Together, these observations strongly emphasise the role of post-transcriptional regulation in 1) selective *VSG* expression; 2) refining the levels of the active-*VSG versus* other expression-site-associated-genes. However, our knowledge of both the mechanisms and molecular players involved remains incomplete.

Here, we used TurboID-mediated proximity labelling combined with quantitative mass spectrometry to identify the protein network involved in post-transcriptional regulation at the ESB. We identified three new ESB components: two ESB-specific, exclusively expressed in the mammalian stage (ESB-specific proteins 2 and 3 – ESB2 / ESB3) and a third that localises to the ESB in bloodstream forms but is also expressed in the insect stage (ESB-associated protein 1 - ESAP1). We show that ESB2 is an RNA nuclease that negatively regulates *ESAG* transcripts. Moreover, mutation of key amino acids required for its nuclease activity significantly impacts its recruitment to the ESB, which is hierarchically dependent on VEX2, ESAP1 and ESB3. Overall, we unravelled a novel molecular mechanism that fine-tunes expression of virulence genes through specialised RNA decay in *T. brucei*.

## Results

### VEX proximity proteomes identify novel components of the ESB and surrounding nuclear bodies

Based on previous studies, we hypothesised that the ESB is likely to have separately assembled ‘transcriptional and processing subdomains’, which are ESB1 and VEX-dependent, respectively [31–33]. To identify the protein network within the ESB ‘processing subdomain’ and at the interface with the SLAB, we performed proximity labelling combined with quantitative mass spectrometry using VEX2 and VEX1 as baits, respectively (**Fig. 1, Supplementary Figs. 1 and 2**). We fused the proteins of interest with TurboID, which labels proteins in living cells and is faster and more efficient than other biotin ligases [40]. In the case of VEX2, given its large size and that its N and C-termini are in closer spatial proximity to the ESB and the SLAB respectively [33], we generated fusions at both termini to increase spatial resolution. We confirmed that VEX1 and VEX2 fusions biotinylated proximal proteins in the expected sub-nuclear compartments upon addition of exogenous biotin (**Supplementary Fig. 1a-e**). Biotinylated material was affinity purified and submitted to mass spectrometry analysis (**Fig. 1a-c, Supplementary Fig. 1f-g**).

**Fig. 1.**
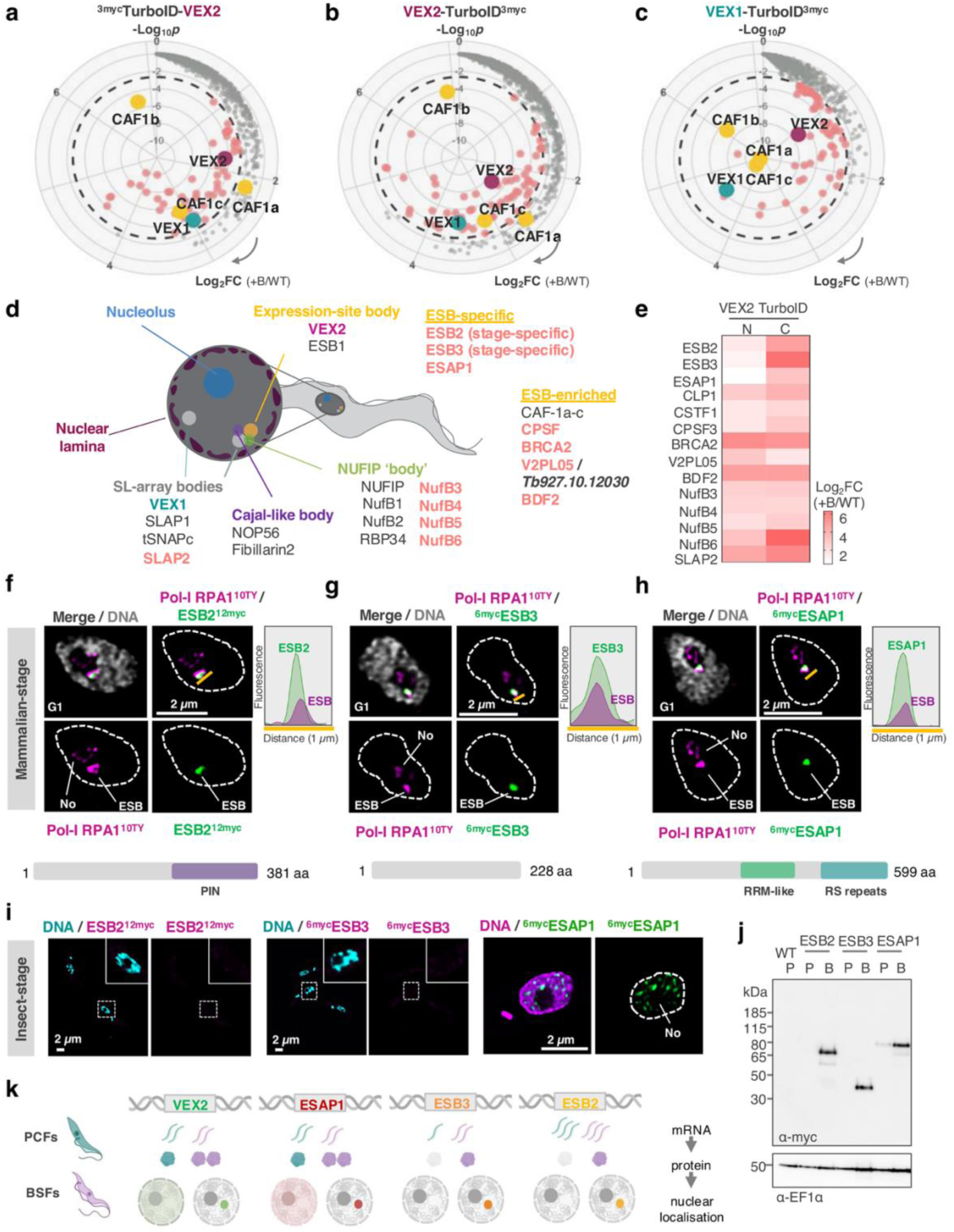
VEX1 and VEX2 TurboID identified novel ESB-specific components, which are developmentally regulated. **a-c**, radial plots representing the proximity proteomes of ^3myc^TurboID-VEX2 (**a**), VEX2-TurboID^3myc^ (**b**) and VEX1-TurboID^3myc^ (**c**). Log_2_FC, which increases clockwise, represents the fold change in protein abundance between the cell lines where VEX1/VEX2 were fused with TurboID and the parental line, both treated with 50 μM of biotin for 18 hours. The y axis (-log_10_*p*) represents statistical significance, which increases towards the centre. VEX1, VEX2 and CAF-1a-c are known interactors, highlighted in cyan, burgundy and yellow, respectively. Datapoints highlighted in salmon represent proteins that are significantly enriched in the TurboID +biotin samples. **d,** cartoon summarising the main nuclear compartments in *T. brucei* bloodstream forms. Known components of each nuclear body (ESB, NUFIP, SLAB or Cajal-like) in grey; new components identified in this study in salmon. **e,** heatmap depicting the Log_2_FC for the newly identified components of the ESB, SLAB and NUFIP body in the ^3myc-TurboID^VEX2 and VEX2^TurboID-3myc^ datasets. **f-h,** fluorescence microscopy analysis of ESB2^12myc^ (**d**), ^6myc^ESB3 (**e**), ^6myc^ESAP1 (**f**) and Pol-I RPA1^10ty^ in bloodstream forms. Histograms on the right-hand side depict the fluorescence signal distribution across the distance highlighted by the yellow lines. A schematic of the number of amino acids, identifiable domains and sequence-motifs is provided for each protein. PIN, RNA nuclease; RRM-like, RNA binding. **i,** fluorescence microscopy analysis of ESB2^12myc^, ^6myc^ESB3 and ^6myc^ESAP1 in procyclic forms. **j,** protein blotting analysis of ESB2^12myc^, ^6myc^ESB3 and ^6myc^ESAP1 expression in bloodstream (B) and procyclic (P) forms. WT, wild-type. EF1α was used as a loading control. The images in **f/g/h/i (right panel)** and **i (left / middle panel)** were acquired using a Zeiss LSM980 Airyscan 2 and a Zeiss AxioObserver, respectively, and correspond to 3D projections by brightest intensity of 0.1 μm stacks. ESB, expression-site body; No, nucleolus. DNA was stained with DAPI (cyan, grey or purple); scale bars: 2 μm. **k,** cartoon summarising how VEX2, ESAP1, ESB3 and ESB2 are developmentally regulated between insect (cyan) and mammalian (purple) stages of *T. brucei* at both RNA and protein levels. In the insect-stage, VEX2 and ESAP1 can be found throughout the nucleus in a speckle-like manner. All proteins form a single protein focus that colocalises with the ESB in the mammalian-form of the parasite. **d/k,** created with BioRender.com

As expected, the baits and known interactors, including the histone chaperone CAF-1a-c [30], were highly enriched in the ‘plus biotin’ samples. In addition and as expected, Pol-I subunits, the Pol-I elongator TDP1 [41], all known telomere binding proteins [20, 42–46] and all known NUFIP body components [36] were also highly enriched. We also detected SIZ1, a known small ubiquitin-like modifier (SUMO) E3 ligase, consistent with the ESB lying within a highly SUMOylated focus [47] (**Supplementary Fig. 2a-c**). Moreover, we identified 49 proteins that had not been previously associated with the ESB or surrounding bodies. GO term analysis revealed that most of them were likely involved in mRNA processing (**Supplementary Fig. 1h-i**), indicating that our efforts to capture the ESB ‘processing subdomain’ were successful. We then ranked 17 of those as ‘high priority hits’ based on 1) fold enrichment in the biotin-treated samples; 2) upregulation in the mammalian stage of the parasite [48]; 3) sequence divergence in African trypanosomes; 4) nuclear speckle-like localisation in insect forms [49] (**Supplementary Fig. 1j**). Among these, 8 were hypothetical proteins of unknown function, which, at first, we designated VEX Proximity Labelling X, V2PLx (*see methods section*). We endogenously tagged the 17 ‘high priority hits’ and assessed their localisation relative to the ESB, SLAB and NUFIP body using fluorescence microscopy.

We identified three novel ESB components, which we designated ESB-specific proteins 2 and 3 (**ESB2**; Tb927.1.1820 / V2PL02 | **ESB3**; Tb927.9.15850 / V2PL01) and ESB-associated protein 1 (**ESAP1**; Tb927.10.11600 / V2PL04) (**Fig. 1d-h; Supplementary Fig. 2d-f**). We found 4 proteins that were accumulated at the ESB, albeit not exclusively: CPSF3, a component of the cleavage and polyadenylation specificity factor [50]; BRCA2, a protein involved in homologous recombination-mediated DNA repair [51]; Tb927.10.12030 (V2PL05), a protein of unknown function; and BDF2, a bromodomain protein involved in epigenetic regulation (**Fig. 1d-e; Supplementary Fig. 3a-i**). Putative components of other assemblies within the cleavage and polyadenylation complex were also significantly enriched in the VEX proximity proteomes: CLP1 (Tb927.11.9760) and CstF1 (Tb927.10.15740) (**Fig. 1e**), which integrate cleavage factor II [52]. Similarly, most of BDF2 known interactors [53] were enriched in the ‘plus biotin’ samples (**Supplementary Fig. 3h**). Notably, BDF2 had been shown to interact with the chromatin at the active-ES [26]. Another bromodomain containing protein, BDF6, and its known interactors, EAF6 (component of the NuA4 histone acetyltransferase complex) and Tb927.10.14190 (V2PL03) [53] were enriched in the VEX proximity proteomes, but all localised to the nucleoplasm with a speckle-like profile without notable enrichment at the ESB (**Supplementary Fig. 3h/j**).

Additionally, we identified a new SLAB component, *SL*-associated protein 2 (SLAP2; Tb927.4.3360 / V2PL07) and 4 new NUFIP body components (NufB3-6), previously identified as KKT (kinetochore components) interacting proteins (KKIP 2/3/4/11) [54] (**Fig. 1d-e; Supplementary Fig. 4**).

We then decided to focus our functional studies on the ESB-specific factors.

### ESB2, ESB3 and ESAP1 are developmentally regulated

ESB2, ESB3 and ESAP1 form a single discrete protein condensate in G1 cells, that colocalises with the ESB in >95% of the nuclei (**Supplementary Fig. 2d-f**), as confirmed by high resolution Airyscan 2 imaging (**Fig. 1f-h**). Structural and sequence-based domain analysis showed that ESB2 contains a C-terminal PIN domain (**Fig. 1f**), which usually functions as a nuclease that cleaves single stranded RNA in a sequence- or -structure-dependent manner [55]. ESAP1 has an RRM-like RNA binding domain and C-terminal arginine/serine (RS) repeats (**Fig. 1h**), both usually present in splicing regulators [56]. No domains could be identified in ESB3 (**Fig. 1g**).

Next, we assessed the expression and localisation of ESB2, ESB3 and ESAP1 in procyclic (insect) forms of *T. brucei* (**Fig. 1i-j, Supplementary Fig. 5**), where *VSG* expression is repressed, and therefore the ESB is absent. RNA-Seq analysis showed that ESB1, ESB2 and ESB3 mRNA levels were modestly upregulated in bloodstream forms, yet no significant changes were observed for VEX1, VEX2 and ESAP1 between the two developmental stages (**Supplementary Fig. 5b**). At the protein level, ESAP1 was downregulated in insect forms and displayed a speckle-like profile throughout the nucleoplasm, whereas ESB2 and ESB3 could not be detected (**Fig. 1i-j, Supplementary Fig. 5a**). These results were also supported by previous comparative transcriptomics and proteomics studies [48, 57] (**Supplementary Fig. 5c-d**). Further, depletion of ESB2 and ESAP1 in procyclic forms using RNAi did not result in any fitness cost or changes to the transcriptome (**Supplementary Fig. 5e-g**). Whilst this result was expected for ESB2, as we showed that it is not expressed in this developmental stage, the function of ESAP1 in procyclic forms remains unclear.

Altogether, the data shows that ESB2 and ESB3 are stage-specific, similarly to ESB1 [31], whereas ESAP1 is expressed in both developmental stages, but ESB-associated in the bloodstream form, similarly to VEX2 [30] – these attributes are reflected in the names that we assigned them. Interestingly, for all known ESB components, the developmental regulation appears to occur mostly at the protein level (**Fig. 1k**).

### ESB2, ESB3 & ESAP1 depletion leads to upregulation of *expression-site associated genes* in bloodstream forms

To study the role of ESB2, ESB3 and ESAP1 in the mammalian-stage, we generated tetracycline inducible RNAi cell lines. Upon induction, we observed a pronounced fitness cost in all three knockdowns, particularly for ESB2 (**Fig. 2a-c**). Successful protein depletion was confirmed by protein blotting (**Fig. 2d-f**). ESB2 always displays two bands, the expected molecular weight aligns with the lower band (59.59 kDa including 12xmyc tag), but the predominant band has a higher molecular weight, suggesting post-translational modifications. Induced and non-induced cells for all three knockdowns homogeneously expressed VSG-2 (the active VSG in our strain) (**Fig. 2g**).

**Figure 2.**
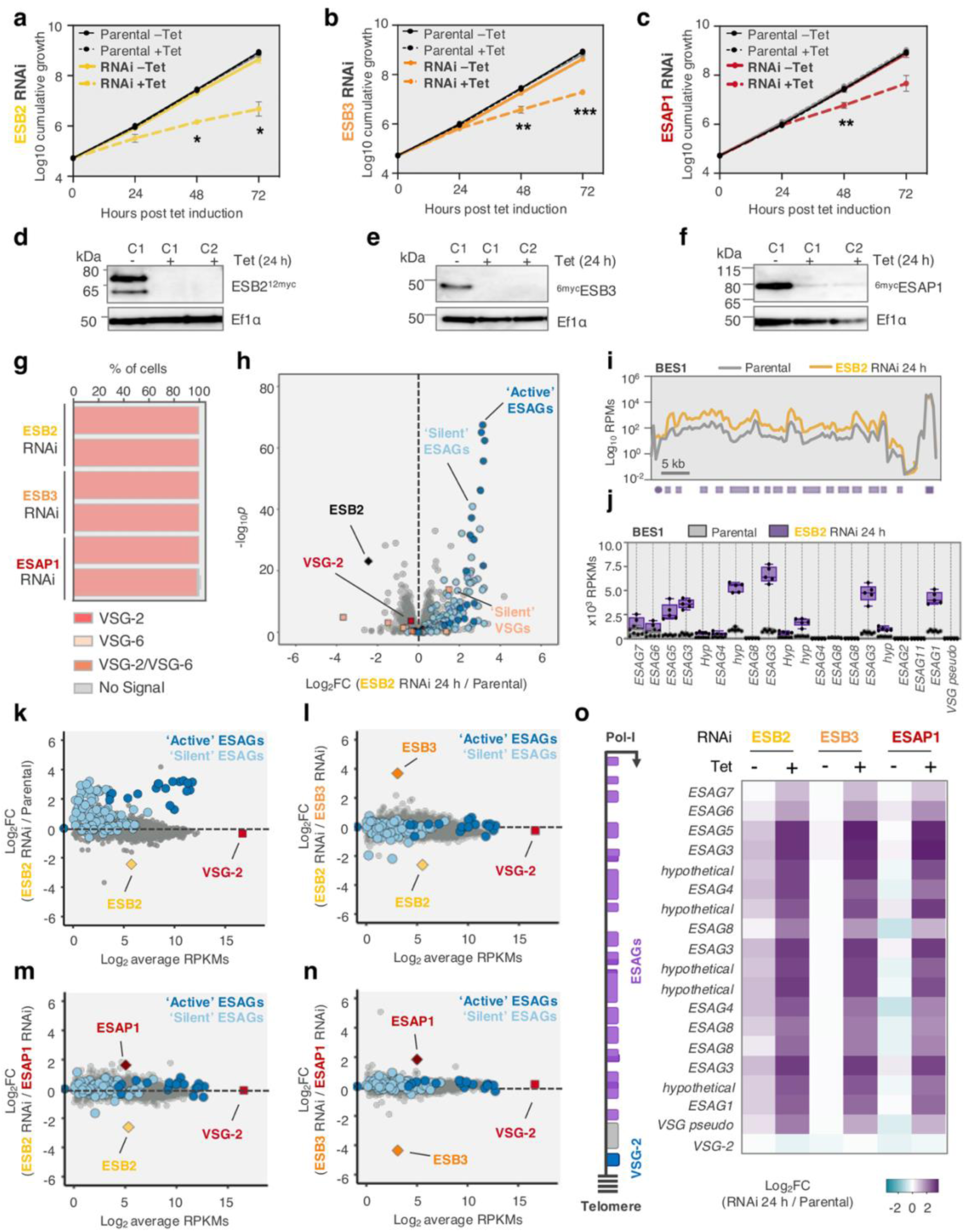
ESB2, ESB3 & ESAP1 knockdown leads to upregulation of *ESAG* transcripts in bloodstream forms. **a-c**, Cumulative growth following tetracycline induced ESB2 (**a**), ESB3 (**b**) and ESAP1 (**c**) knockdown; \**p* < 0.05; \*\**p* < 0.01, \*\**p* < 0.001 (multiple *t* tests). **d-f,** protein blotting analysis of ESB2 (**d**), ESB3 (**e**) and ESAP1 (**f**) expression following their respective knockdowns. C1, clone 1; C2, clone 2. EF1α, loading-control. **g,** immunofluorescence analysis of VSG expression following ESB2/ESB3/ESAP1-knockdown (48 h); >100 cells per condition were analysed. **a-c/g,** error bars (not visible in **g**), SD; data are averages from two independent biological replicates and representative of independent experiments. **h-o,** RNA-Seq analysis of ESB2 (n=5), ESB3 (n=3) and ESAP1 (n=3) knockdowns at 24 hours post induction, where *n* values correspond to the number of biological replicates. **h,** volcano plot depicting upregulation of ‘active’ and ‘silent’ *ESAGs* following ESB2 RNAi. **i,** transcript abundance at the active *VSG-2* locus. Purple circle, promoter; purple box, *VSG-2*; lilac boxes, *ESAGs*. RPM, reads per million. Bin size 0.5 kb. **j,** box plots depicting *ESAG* transcript abundance in RPKMs (reads per kilobase per million) at the active-ES in the parental line *versus* ESB2 RNAi. Boxes span between the 25^th^ and 75^th^ percentile, the line corresponds to the mean value; whiskers span between minimum and maximum value; all datapoints are depicted. hyp, hypothetical. **k-n,** scatter plots comparing average gene expression in the parental line (in RPKMs) with fold change in expression in ESB2 RNAi 24 h / Parental (**k**), ESB2 RNAi 24 h / ESB3 RNAi 24 h (**l**), ESB2 RNAi 24 h / ESAP1 RNAi 24 h (**m**) and ESB3 RNAi 24 h / ESAP1 RNAi 24 h (**n**). **o,** heatmap depicting the Log_2_FC within the active-ES for induced and uninduced samples of ESB2, ESB3 and ESAP1 RNAi normalised against the parental line. ESB2 RNAi non-induced samples present some leaky expression. From top to bottom, genes are ordered as they appear in BES1 (active-ES). *ESAG2* and *ESAG11* were omitted as their expression could not be detected in all replicates in the parental line when applying high stringency mapping.

We then conducted RNA-Seq analysis at 24 hours post induction (**Fig. 2h-o, Supplementary Fig. 6**). ESB2, ESB3 and ESAP1 depletion led to a striking and specific upregulation of all *ESAG* transcripts from the active *VSG*-ES (up to 11-fold), accompanied by a modest reduction in the active-*VSG* transcript abundance. Additionally, we also observed a significant increase in some *ESAG* transcripts originating from ‘silent’ *VSG*-ESs, without any major changes in the mRNA levels of the corresponding *VSGs* (**Fig. 2h/k, Supplementary Fig. 6**).

This hinted that ESB2, ESB3 and ESAP1 somehow negatively regulate the expression of *ESAGs*. Further, the transcriptional profiles following the knockdown of either of these factors were remarkably similar (**Fig. 2l-o, Supplementary Fig. 6**), suggesting that they might operate in the same pathway. To test this hypothesis, we proceeded to investigate the co-dependency between ESB2 / ESB3 / ESAP1 and other ESB-associated factors.

### ESB2 recruitment hierarchically depends on VEX2, ESAP1 and ESB3

Firstly, we focused on the ‘interactions’ between ESB2 / ESB3 / ESAP1 and assessed how they reciprocally impacted their relative mRNA and protein abundances as well as localisation to the ESB (**Fig. 3a-c, Supplementary Fig. 7a-e**). ESAP1 localisation was not affected by ESB2 or ESB3 knockdown and was therefore ESB2 and ESB3-independent. Its protein expression was also not significantly affected by VEX2, ESB2 or ESB3 knockdowns (**Fig. 3b-c, Supplementary Fig. 7a/c**). Next, ESB3 localisation was shown to be ESAP1-dependent, with a significant reduction in the percentage of nuclei where an ESB3 focus could be detected following ESAP1 RNAi, but without significant changes in protein abundance. Further, while ESB3 localisation was ESB2-independent, its protein levels were significantly reduced (**Fig. 3b-c, Supplementary Fig. 7a/d**). Lastly, we observed that ESB2 localisation depended on both ESAP1 and ESB3; in uninduced cells, ESB2 foci could be detected in ∼97.4% of nuclei, whereas following both knockdowns, the foci could no longer be detected in ∼78% of nuclei. Additionally, in the absence of ESB3, ESB2 protein levels were significantly reduced (**Fig. 3a-c, Supplementary Fig. 7e**). Interestingly, ESB2 and ESB3 reciprocally modulate each other’s protein levels (**Fig. 3c**); transcriptomic data showed no significant changes at the mRNA level (**Supplementary Fig. 7b**).

**Figure 3.**
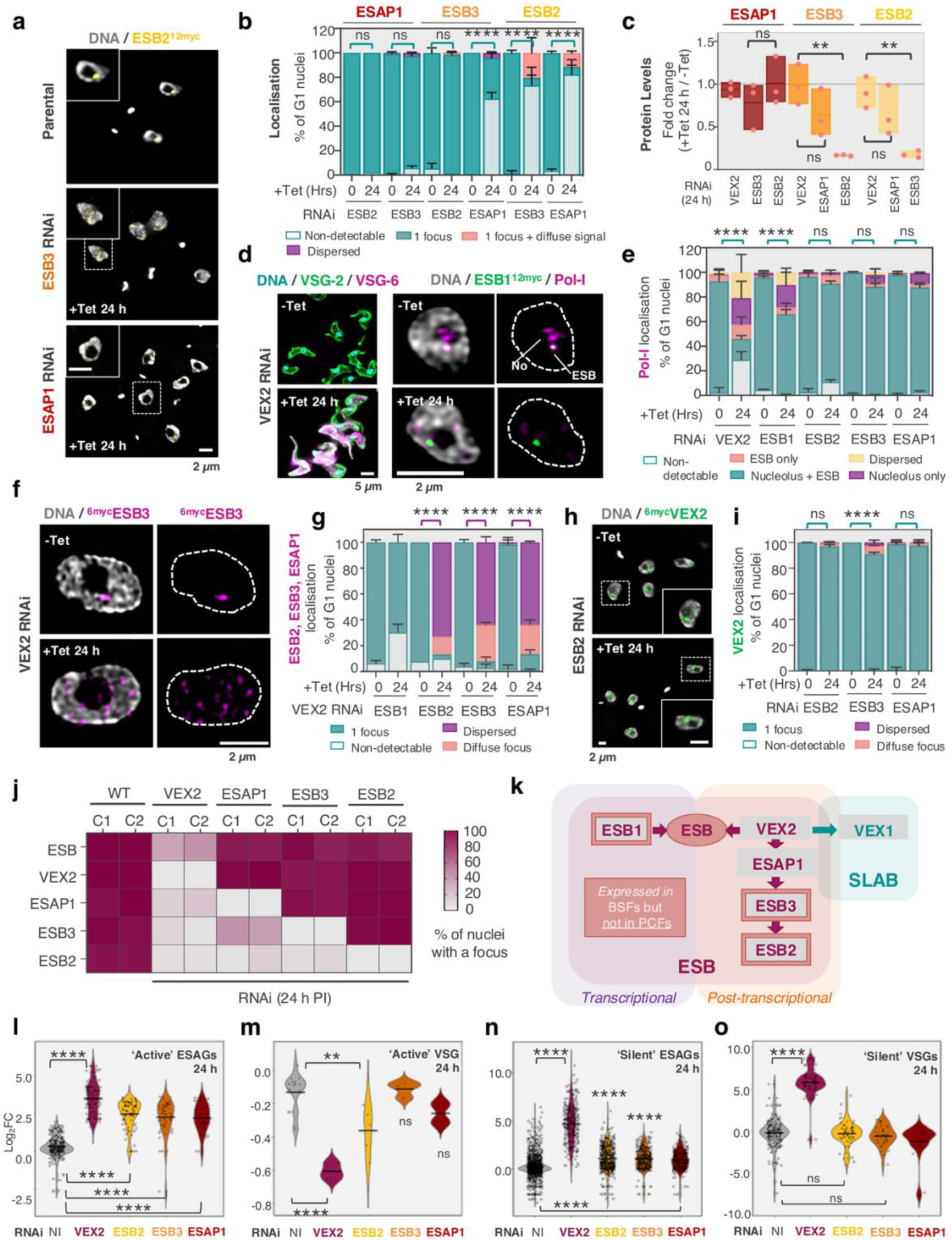
A hierarchy of co-dependencies at the ESB negatively regulates *ESAGs* and silent *VSGs* expression in a ESB2 and VEX2-dependent manner, respectively. **a-b**, Fluorescence microscopy analysis of ESB2^12myc^ localisation following ESB3 and ESAP1 RNAi (**a/b**), ^6myc^ESB3 localisation following ESB2 and ESAP1 RNAi (**b**) and ^6myc^ESAP1 localisation following ESB2 and ESB3 RNAi (**b**). **c,** ESB2, ESB3 and ESAP1 protein levels following VEX2, ESB2, ESB3 and ESAP1 RNAi. The values were derived from protein blotting analysis of three biological replicates per cell line. Two-tailed paired Student’s *t* tests were applied; ns, non-significant; \*\**p* < 0.01. Fluorescence microscopy analysis of Pol-I localisation following VEX2, ESB1, ESB2, ESB3 and ESAP1 knockdown (**d-e**); ESB1^12myc^, ESB2^12myc^, ^6myc^ESB3 and ^6myc^ESAP1 localisation following VEX2 knockdown (**f-g**); ^6myc^VEX2 localisation following ESB2, ESB3 and ESAP1 knockdown (**h-i**). The images in **a/d (left)/h** were acquired using a Zeiss AxioObserver, whereas the images in **d(right)/f** were acquired using a Zeiss LSM980 Airyscan 2. All correspond to 3D projections by brightest intensity of 0.1 μm stacks. DNA was stained with DAPI (cyan or grey); scale bars: 2 or 5 μm. The graphs in **b/e/g/i** depict averages of two biological replicates; >100 G1 cells per condition were analysed; all analyses were performed at 24 hours post induction. Two-way ANOVA; ns, non-significant; **** *p*<0.0001. The colour of the lines below the statistical results indicates which categories (1 focus, cyan; dispersed, purple) were compared between conditions. **j,** heatmap summarising the microscopy analyses; C1, clone 1; C2, clone 2. **k,** cartoon depicting the complex network of co-dependencies at the ESB and interface with the SLAB. BSFs, bloodstream forms; PCFs, procyclic forms. The direction of the arrows indicates that the protein upstream is required for the localisation of the protein downstream to its corresponding nuclear compartment. **l-o,** violin plots depicting a comparative transcriptomic analysis between VEX2, ESB2, ESB3 and ESAP1 RNAi cell lines with a focus on Pol-I transcribed gene cohorts: ‘active’ *ESAGs* (**l**), ‘active’ *VSG* (**m**), ‘silent’ *ESAGs* (**n**) and ‘silent’ *VSGs* (**o**). Log_2_FC, fold change in transcript abundance between RNAi (uninduced or 24 hours post induction) and the parental cell line. The violins span between minimum and maximum values, centre lines correspond to the mean, all datapoints are shown. One-way ANOVA; ns, non-significant; ** *p*<0.01; **** *p*<0.0001.

Secondly, we investigated whether the ESB integrity was impacted by ESB2, ESB3 and/or ESAP1 depletion; we used VEX2 and ESB1 knockdowns as controls (**Fig. 3d-e, Supplementary Fig. 7f**). ESB1 knockdown reduced the percentage of nuclei where an ESB could be detected by approximately 22% (**Fig. 3e**). VEX2 depletion, which disrupts *VSG* monogenic expression, resulting in >90% of cells expressing at least two VSGs, had a much more significant impact on Pol-I localisation. An ESB could only be detected in 28.6% of nuclei, and the nucleolar pool of Pol-I was also significantly affected (**Fig. 3d-e**). These results were consistent with previous reports [30, 31]. In contrast, ESB2, ESB3 and ESAP1 depletion did not significantly impact the ESB integrity (**Fig. 3e, Supplementary Fig. 7f**).

Thirdly, since 1) VEX2 has a major role in maintaining, possibly establishing the ESB structure; 2) ESB2, ESB3 and ESAP1 were found in proximity to VEX2; we next asked whether their localisation was VEX2-dependent. VEX2 knockdown led to a pronounced nuclear dispersion of ESB2, ESB3 and ESAP1 in >64% of nuclei (**Fig. 3f-g, Supplementary Fig. 7g**). Conversely, ESB2, ESB3 and ESAP1 RNAi did not impact VEX2 localisation (**Fig. 3h-i, Supplementary Fig. 7h**).

Altogether, these observations allowed us to define a complex network of co-dependencies at the ESB (**Fig. 3j/k**), whereby its integrity depends on both VEX2 and ESB1, albeit by independent mechanisms. Additionally, VEX2 liaises the ESB to the SLAB via VEX1 [32], and is required for the sequential recruitment of ESAP1, ESB3 and ESB2 to the ESB.

We then performed a comparative transcriptomic analysis (**Fig. 3l-o, Supplementary Fig. 7i-k**) using the RNA-Seq data generated for ESB2, ESB3 and ESAP1 RNAi cell lines in this study and previously published RNA-Seq data for VEX2 RNAi [30]. The most significant changes following VEX2 depletion were the upregulation of ‘silent’ *ESAGs* (32-fold) and ‘silent’ *VSGs* (dual promoter 344-fold; single promoter 42-fold), which reflect activation of silent *VSG*-ESs. ‘Silent’ *ESAGs* were also upregulated following ESB2, ESB3 and ESAP1 RNAi, albeit to a lesser extent (**Fig. 3n-o**). Other Pol-I transcribed transcripts (*mVSGs, procyclins, PAGs*) were upregulated in all conditions, but most significantly following VEX2 depletion (**Supplementary Fig. 7i-k**). Notably, the most significant change following ESB2, ESB3 and ESAP1 knockdown was the upregulation of ‘active’ *ESAGs* (**Fig. 3l**). Considering the hierarchy of co-dependencies we established (**Fig. 3k**), the modulation of ‘active’ *ESAGs’* mRNA abundance is likely to be mediated by ESB2; similar changes observed in the VEX2, ESB3 and ESAP1 RNAi samples are possibly indirect and cannot be differentiated from the effect of ESB2 mislocalisation and/or downregulation. Thus, we then focused our subsequent efforts on ESB2.

### ESB2 is an RNA nuclease and inactivation of key catalytic residues impacts recruitment to the ESB

Based on protein sequence, ESB2 has orthologues in other trypanosomes and *Leishmania*, whereas ESAP1 has no orthologues beyond trypanosome species. Both ESB2 and ESAP1 share the highest sequence similarity with orthologues in *T. b. gambiense, T. equiperdum and T. evansi*, which are the only species where an orthologue for ESB3 can be found, and the only ones that possess *ESAGs* among trypanosome species (**Supplementary Fig. 8a-d**). However, when inspecting ESB2’s PIN domain, we found that the closest experimentally determined structural homologue occurred in human SMG6 (**Fig. 4a-c**), an endonuclease and critical component of the nonsense-mediated decay (NMD) pathway, which is mostly known for the degradation of aberrant transcripts [58]. Notably, the catalytic residues required for the nuclease activity are strictly conserved (**Fig. 4c, Supplementary Fig. 9**).

**Figure 4.**
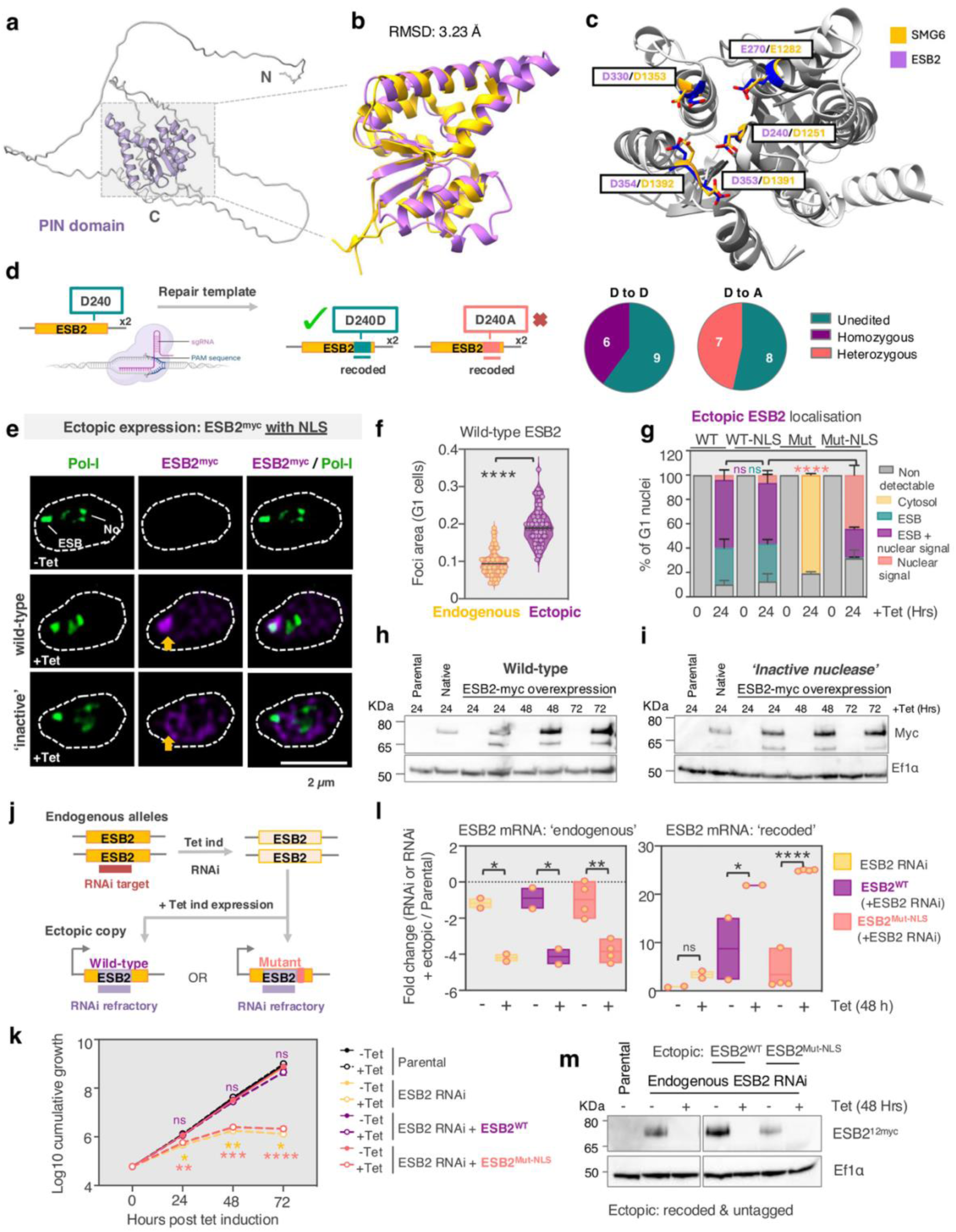
ESB2 localisation and function are dependent on its RNA nuclease activity. **a-c**,ESB2 PIN domain analysis. **a**, AlphaFold2 prediction of ESB2 structure (PIN domain highlighted in purple). **b-c**, predicted ESB2 PIN domain superimposed with the PIN domain structure of human SMG6 (PDB ID 2HWW). Root mean square deviation (RMSD) quantifies the average distance between corresponding atoms in two superimposed structures. Catalytic residues are highlighted in **c**. **d,** CRISPR/Cas9 mediated precision editing of ESB2 D240 to alanine. No homozygous clones for the desired mutation could be generated, however successful double allele editing was achieved when using a repair template containing a synonymous mutation. 15 clones were screened for both synonymous and non-synonymous mutations, respectively. The data are representative of two independent experiments. **e-i,** tetracycline inducible overexpression of ESB2^12myc^ in bloodstream forms. **e-g,** fluorescence microscopy analysis of Pol-I and ectopically expressed ESB2^12myc^. Wild-type or catalytic mutant (D240A, D330A, D353A), with or without an La-NLS sequence, were analysed. Images were acquired using a Zeiss LSM980 Airyscan 2 and correspond to 3D projections by brightest intensity of 0.1 μm stacks. DNA was stained with DAPI (grey); scale bars: 2 μm. The violin plot (**e**) displays data from >100 G1 cells and is representative of two biological replicates and independent experiments. The stacked graph (**f**) depicts averages of two biological replicates; >100 G1 cells per condition were analysed; error bars indicate standard deviation. The statistics depicted on the graph are focused on % of cells with an ESB (cyan), ESB + nucleolar signal (purple) and nuclear signal (salmon). Data in **e-g** corresponds to 24 hours post induction. **h-i,** protein blotting analysis of wild-type (**h**) or mutant (**i**) ESB2 expression. ‘Native’ corresponds to a cell line where ESB2 was endogenously tagged. **j-m,** tetracycline inducible expression of ESB2^Mut-NLS^ and ESB2^WT^ upon depletion of endogenous ESB2. **j**, schematics summarising the cell lines that were generated. The ectopic copies were recoded to be refractory to the RNAi target sequence and were untagged. **k**, cumulative growth following tetracycline induction. The ESB2 RNAi cell line was used as a benchmark. Regarding the statistical analysis, the colours indicate which cell line they refer to and correspond to induced versus non-induced conditions for each timepoint. Error bars (not visible) correspond to SD. **l**, RT-qPCR analysis showing successful expression of *ESB2^Mut-NLS^* and *ESB2^WT^* ectopic copies upon endogenous ESB2 depletion. Primers annealed in the *ESB2* 5’UTR and beginning of its coding sequence amplifying only endogenously expressed *ESB2* (left hand side) or in the recoded region amplifying only ectopically expressed *ESB2* (right hand side). The expression of the gene of interest was normalised against actin (housekeeper gene) and displayed as fold change in relation to the parental cell line. Each experiment was conducted using three technical replicates per condition, which were averaged. At least two independent clones were tested (all datapoints are displayed). **m,** protein blotting analysis showing the depletion of endogenous ESB2, which was C-terminally tagged with 12xmyc in the presence of ectopically expressed *ESB2^Mut-NLS^* or *ESB2^WT^*. Statistical analysis was performed as follows: two-tailed unpaired Student’s *t* test (**f**); two-way ANOVA (**g**); multiple two-tailed paired Student’s *t* tests (**k**); one-way ANOVA (**l**). ns, non-significant; ** *p* < 0.01; *** *p* < 0.001; **** *p* < 0.0001. **h, i, m,** EF1α was used as loading-control. The violin plot (**f**) and the floating bar graphs (**l**) span between minimum and maximum values, centre lines correspond to the mean, all datapoints are shown.

We used CRISPR/Cas9-mediated precision editing to introduce a point mutation in D240 (D1251 in humans), which has been shown to render the nuclease inactive [59]. While we obtained transfectants where both alleles contained a synonymous mutation, we were unable to generate mutants where D240 had been replaced by an alanine in both alleles (**Fig. 4d, Supplementary Fig. 8e-g**). Based on the fitness cost caused by its depletion, ESB2 is likely to be essential for parasite survival in a nuclease-dependent manner.

Since we could not generate viable mutants when editing the endogenous alleles, we then decided to express a tetracycline inducible ectopic copy of ESB2, either wild-type or a catalytic mutant (**Fig. 4e-i, Supplementary Fig. 10a-b**). Depending on the PIN domain, one to three of the catalytic residues may need to be mutated to completely inactivate the nuclease [59, 60]. To be sure, we mutated three aspartic acids (D240A, D330A, D353A), as previously performed for human SMG6 [61]. First, we assessed protein localisation. Overexpression of wild-type ESB2 led to its accumulation at the ESB in 85.8% of nuclei often with additional nucleoplasmic signal; the focus at the ESB was larger than the one observed when the protein was expressed at endogenous levels. To our surprise, ESB2 containing an inactive nuclease, when detected, localised solely to the cytosol (**Fig. 4e-g, Supplementary Fig. 10a**). Notably, the triple replacement abrogates the nuclease activity and/or RNA-binding capacity but does not compromise protein folding or stability [61, 62]. It is conceivable that somehow these mutations led to a conformational change that either obstructed the nuclear localisation signal (NLS) or ESB2’s ability to interact with a binding partner that transports it into the nucleus through a piggyback mechanism. We then decided to fuse this mutant version of ESB2 with a known NLS to drive it into the nucleus. The mutant could then be detected in 68.4% of nuclei, but only in 24.2% of which, it clearly accumulated at the ESB by microscopy. A wild-type version fused with the same NLS was generated as a control, and no changes to its localisation were observed (**Fig. 4e/g**). Protein blotting analysis showed that both wild-type and mutant versions of ESB2 were expressed at similar levels (**Fig. 4h-i**); neither had an impact on the percentage of cells where an ESB could be identified or its morphology (**Supplementary Fig. 10b**).

Next, we asked whether ectopic expression of wild-type or mutant ESB2 could rescue the RNAi-mediated loss of fitness. To this end, we engineered constructs with dual-inducible ESB2 RNAi and overexpression. The overexpression sequences, both wild-type and mutant were recoded so that they were refractory to the RNAi targeting sequence, which would then only target endogenous *ESB2* transcripts (**Fig. 4j**). Ectopic copies were untagged and in the case of the mutant, we fused it with an NLS to ensure that it reached the nucleus. The ectopic wild-type copy of ESB2 could rescue the RNAi phenotype, but not the catalytic mutant (**Fig. 4k**). Depletion of endogenous *ESB2* and expression of the ectopic copies was confirmed at the RNA level (**Fig. 4l**); depletion of endogenous ESB2 was also confirmed at the protein level (**Fig. 4m**).

Collectively, these experiments show that ESB2’s nuclease activity (and/or RNA binding capacity) is required for function and recruitment to the ESB.

### ESB2 negatively regulates *ESAG* transcripts in a nuclease-dependent manner

We then ectopically expressed tetracycline-inducible untagged versions of either wild-type or mutant ESB2 (**Fig. 5a-b**) and analysed their transcriptomes. We observed that overexpression of wild-type or catalytically inactive ESB2 led to a downregulation or upregulation of *ESAGs* at the active-*VSG*-ES, respectively (**Fig. 5c-d, Supplementary Fig. 10c**). Notably, the transcriptional profile obtained following overexpression of the catalytic mutant resembled that of the ESB2 knockdown (**Fig. 5e, Supplementary Fig. 10d**), suggesting that the mutant competes with the endogenous wild-type protein at the ESB. The phenotype was less penetrant than the RNAi, presumably given the mutant’s poor recruitment to the ESB (**Fig. 4e/g**).

**Figure 5.**
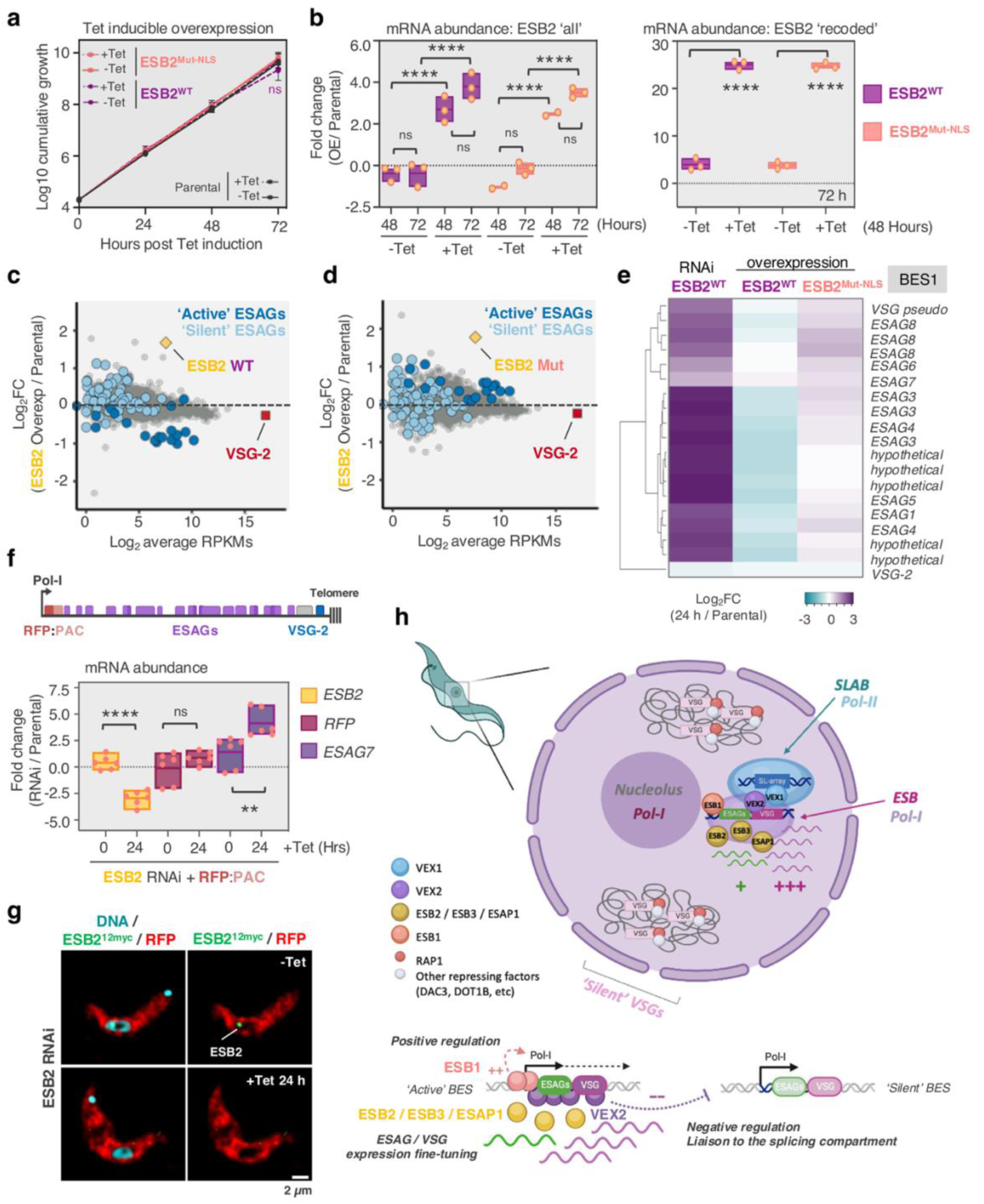
ESB2 negatively regulates *ESAG* transcripts in a nuclease activity-dependent manner. **a-e**, tetracycline inducible overexpression of untagged ESB2 in bloodstream forms. **a**, cumulative growth following tetracycline induced ESB2^Mut-NLS^ (salmon), and ESB2^WT^ (purple) expression; ns, non-significant (multiple *t* tests). Error bars indicate standard deviation. **b**, determination of *ESB2* mRNA levels by RT-qPCR. The primers annealed outside the recoded region amplifying both endogenously and ectopically expressed *ESB2* (left hand side) or in the recoded region amplifying only ectopically expressed *ESB2* (right hand side). One-way ANOVA; ns, non-significant; **** *p*<0.0001. **c-e,** RNA-Seq analysis of ESB2^WT^ and ESB2^Mut-NLS^ overexpression at 72 hours post induction. Three biological replicates were used as well as parental and uninduced controls. **c-d,** scatter plots comparing average gene expression in the parental line (in RPKMs, reads per kilobase per million) with fold change in expression in ESB2^WT^ overexpression over parental (**c**) and ESB2^Mut-NLS^ overexpression over parental (**d**). **e,** heatmap depicting the Log_2_FC within the active-ES (BES1) for induced samples of ESB2^WT^ and ESB2^Mut-NLS^ overexpression compared to ESB2 RNAi – all normalised against the parental line. Hierarchical clustering – average linkage, Euclidean distance. *ESAG2* and *ESAG11* were omitted as their expression could not be detected in all replicates in the parental line when applying high stringency mapping. **f-g**, analysis of cell lines containing an RFP:PAC fluorescent reporter at the active *VSG*-ES before and after ESB2 RNAi. **f**, determination of *RFP*, *ESAG7* and *ESB2* mRNA levels by RT-qPCR. Two-tailed paired Student’s *t* test; ns, non-significant; ** *p*<0.01; **** *p*<0.0001. **i**, fluorescence microscopy analysis. DNA was stained with DAPI (cyan); scale bars: 2 μm. For all RT-qPCR experiments (**b/f**), the expression of the gene of interest was normalised against actin (housekeeper gene) and displayed as fold change in relation to the parental cell line. Each experiment was conducted using three technical replicates per condition, which were averaged. Three independent clones of ESB2 overexpression or RNAi were tested; in **f**, two independent experiments are depicted. The floating bar graphs span between minimum and maximum values, centre lines correspond to the mean, all datapoints are displayed. **h,** cartoon depicting the nuclear architecture of *T. brucei* bloodstream forms, including all known components of the ESB and the VEX-mediated interaction with one of the *SL*-arrays. ESB1 is a transcriptional activator required for transcription at the active-ES. VEX2 is an exclusion factor, which prevents activation of silent *VSGs* and therefore a negative regulator. ESAP1 and ESB3 are required for ESB2 recruitment, which ultimately negatively regulates *ESAGs* expression. Created with BioRender.com

Lastly, we wondered if ESB2’s role in modulating the expression of *ESAGs* was specific to this gene cohort or linked to the position within the transcription-unit. To test this, we used an RFP:PAC reporter under the control of tubulin 5’ and 3’UTRs (unrelated DNA elements), which was integrated immediately downstream of the active-ES promoter and therefore immediately upstream of the first *ESAG* gene, *ESAG7* (**Fig. 5f-g**). We found no significant differences in *RFP* expression between the parental line and induced and uninduced samples of ESB2 RNAi. As expected, *ESB2* and *ESAG7* transcripts were down- and upregulated following ESB2 RNAi, respectively (**Fig. 5f**). ESB2 depletion and RFP expression were also confirmed at the protein level (**Fig. 5g**). We cannot exclude that additional layers of post-transcriptional regulation of *RFP* transcripts might bypass changes at the ESB, however, these results strongly suggest that there is specificity for *ESAG* transcripts.

In summary, ESB2 negatively regulates *ESAG* transcripts in a nuclease dependent manner.

## Discussion

To this day, there are significant gaps in our understanding of the mechanisms underpinning single gene choice and antigenic switching, including those governing antigenic variation in parasites that cause malaria and sleeping sickness [3, 4]. In *T. brucei*, the ESB accommodates the single active*-VSG-*ES, which not only encodes for the active-*VSG*, but also other genes involved in key host-parasite interactions.

The reason why *ESAG*s are present in *VSG*-ESs has remained a mystery. A plausible explanation is that this location confers mammalian-specific expression, especially considering that trypanosomes have limited control over transcription initiation in Pol-II transcription-units [63]. Another prevailing theory is that *ESAGs* are involved in host adaptation. For instance, trypanosomes, like many other pathogens, scavenge iron from their mammalian host through transferrin uptake. Their transferrin receptors are heterodimers of ESAG6 and ESAG7, whereby only one receptor is expressed at a time. Whilst sequence variation was originally thought to enable uptake of transferrin from different mammals [9, 64], recent structural and biochemical studies elegantly showed that polymorphic sites are mostly found in exposed regions not involved in transferrin binding, suggesting that receptor diversification might be driven by a need for antigenic variation to prolong survival in the host [19]. Whether other *ESAGs* are involved in host adaptation and/or contribute to antigenic variation is yet to be determined.

Overall, despite their critical roles, *ESAG* gene products are required at much lower levels than the active-VSG, which forms a dense coat on the trypanosome surface. Indeed, despite being co-transcribed in the same polycistronic transcription-unit, under the same Pol-I promoter, *ESAGs* are expressed at significantly lower levels. While several post-transcriptional mechanisms have been implicated in enhancing *VSG* mRNA processing and stability [32, 33, 36–38], no mechanism had been previously linked to the regulation of *ESAGs*, although a recent genome wide untranslated region (UTR) screen suggested *ESAGs* UTRs might contain negative regulatory sequences [65]. Here, we identified the molecular machinery that negatively regulates *ESAG* transcripts (**Fig. 5h, Supplementary Fig. 10e**). Combined, these different processes lead to a differential expression of >140-fold between *ESAGs* and the active-*VSG*.

Notably, twenty-four years after its discovery, only two ESB components had been identified: ESB1 and VEX2 [30, 31]. Here, we identified three novel factors, ESB2, ESB3 and ESAP1, and more importantly, we identified a novel layer of post-transcriptional regulation within the ESB. These factors are developmentally regulated, raising questions as to how their expression is repressed in the insect-stage and subsequently reactivated. Additionally, we found a complex network of co-dependencies that supports our initial prediction that the ESB has separately assembled subdomains, associated with transcriptional and post-transcriptional regulation. ESB1 is a transcriptional activator enriched near the Pol I promoter [31], whereas VEX2 is a multi-functional factor with a central role in maintaining ESB integrity [30], preventing activation of silent *VSG*-ESs [30, 33] and coordinating post-transcriptional regulatory mechanisms. The latter include: 1) enhancing *VSG* mRNA processing through spatial proximity to the *SL*-array via VEX1 [32, 33]; and 2) assembling the ESAP1 / ESB3 / ESB2 axis for negative regulation of *ESAGs* expression. Through VEX2’s proximity proteome, we also identified new components of the SLAB, involved in the production of *SL*-RNA; new components of the NUFIP body, likely involved in the production of splicing components and/or *SL*-RNA modification [36]; and a remarkable enrichment of CPSF3, a key component of the CPSF complex [66] at the ESB. This restricted access to mRNA processing machinery aligns with previous reports that the low amounts of transcripts generated from silent *VSG*-ESs are not efficiently polyadenylated [67]. These observations are consistent with a role for VEX2 in the spatial integration of *VSG* transcription and RNA processing in the 3D nuclear space [32, 33].

Interestingly, while NMD is dispensable in *T. brucei* [68–71] and none of VEX2, ESAP1, ESB3 or ESB2 are homologues of its major regulators, they do share structural features with some of its components. VEX2 has an Up-frameshift-1 (UPF1)-like helicase domain [30], ESAP1 has an UPF3-like RRM domain and ESB2 has an SMG6-like PIN domain (**Supplementary Fig. 9**). NMD is widely known as an RNA surveillance pathway responsible for the degradation of aberrant transcripts in metazoa, such as those containing premature stop codons. However, it has been increasingly associated with gene expression regulation to facilitate cellular responses to environmental changes and genome maintenance. In fact, in mammals, ∼10% of its target transcripts are unmutated [72, 73] and recent studies have showed that UPF and SMG proteins, originally discovered to function in NMD, also have roles in other pathways. For instance, UPF1 has been implicated in telomere homeostasis [74, 75] and specialised mRNA decay pathways, including regulated degradation of replication-dependent histone mRNAs [75, 76]. SMG6 has also been shown to be enriched at the telomeres and negatively regulate TERRA association with chromatin in mammals [77, 78].

More broadly, in bacteria and archaea, PIN domains are primarily associated with toxin–antitoxin systems [79]. In eukaryotes, they are usually found as parts of larger proteins and/or protein complexes, involved in a range of processes involving RNA cleavage, including not only NMD but also ribosomal RNA biogenesis, RNA interference and exosome-mediated RNA degradation [62]. While some nucleases can be highly specific to a single RNA target, others are sequence independent [62, 80]. However, the fundamental principles of substrate recognition and cleavage by PIN containing proteins remain elusive, mostly challenged by the difficulty in stabilising the protein-RNA complex for structural studies [62].

Therefore, the mechanism underpinning ESB2-mediated *ESAG* transcript regulation remains to be determined. Since eukaryotic PIN domains tend to be part of larger protein complexes, it is conceivable that the interaction with a binding partner could either direct substrate specificity and/or activate the nuclease that is otherwise kept in an ‘off state’. It is tempting to speculate that ESB3 could be one such interacting partner, given that orthologues can only be found in trypanosomes that possess *ESAGs* and the fact that ESB2/ESB3 modulate each other’s abundance. Further, ESB2 displays a higher molecular weight than predicted, consistent with the presence of post-translational modifications, which remain undefined. Several ESB components are likely to be SUMOylated and/or contain SUMO-interacting motifs (SIMs) [47] – based on *in silico* prediction [81], ESB2 contains a SIM. SUMOylation has been previously implicated in regulating the stability and function of several nucleases [82, 83]. Additionally, some SMG factors are regulated through phosphorylation [72].

In summary, we identified a novel post-transcriptional mechanism of gene expression control, which is spatially regulated at a sub-telomeric location, within a dedicated nuclear body, and fine-tunes expression of proteins involved in host-pathogen interactions and immune evasion in *T. brucei*. Notably, if we consider toxin-antitoxin systems in bacteria and that an exonuclease is also responsible for silencing genes linked to severe malaria [84], nucleases may indeed play a broader role in regulating pathogen-specific gene expression programmes. Overall, our work underscores how specialised RNA decay can regulate expression of specific genes.

## Methods

### *T. brucei* growth and manipulation

Bloodstream-form *T. brucei*, Lister 427 (L427), 2T1 cells [85] and 2T1/T7/Cas9 cells [86] were grown in HMI-11 medium; procyclic-form *T. brucei*, Lister 427 (L427) and PT1 cells were grown in SDM-79 medium; and genetically manipulated using electroporation [87]; cytomix or human T cell nucleofector solution (Lonza) were used for all transfections.

In bloodstream forms, puromycin, phleomycin, hygromycin, neomycin and blasticidin were used at 2, 2, 2.5, 2 and 10 µg ml^−1^ for selection of recombinant clones; and at 1, 1, 1, 1 and 2 µg ml^−1^ for maintaining those clones, respectively. Cumulative growth curves were generated from cultures seeded at 10^5^ cells ml^−1^, counted on a haemocytometer and diluted back to 10^5^ cells ml^−1^ as necessary.

In procyclic forms, puromycin, phleomycin, hygromycin, neomycin and blasticidin were used at 2, 2.5, 50, 10 and 10 µg ml^−1^ for selection of recombinant clones; and at 1, 1, 1, 1 and 2 µg ml^−1^ for maintaining those clones, respectively. Cumulative growth curves were generated from cultures seeded at 10^6^ cells ml^−1^, counted on a haemocytometer and diluted back to 10^6^ cells ml^−1^ as necessary.

Tetracycline was applied at 1 µg ml^−1^ for RNAi, overexpression or Cas9 induction.

### Plasmids and constructs

For epitope-tagging at the native locus [88], a previously generated plasmid was used: pNAT^TAGx^ to add an N-terminal 6× c-myc to VEX2 [30] and pNAT^xTAG^ to add an C-terminal 12× c-myc to BRCA2 (Tb927.1.640) [51]. Additionally, twenty-four new constructs were generated for the present study. pNAT^xTAG^ to add a C-terminal 12× c-myc to ESB1 (Tb927.10.3800), BDF6 (Tb927.1.3400), EAF6 (Tb927.9.2910), V2PL02 / ESB2 (Tb927.1.1820), V2PL03 (Tb927.10.14190), V2PL05 (Tb927.10.12030), V2PL06 (Tb927.10.15870), V2PL07 / SLAP2 (Tb927.4.3360), V2PL08 (Tb927.10.11230), KKIP2 / NufB3 (Tb927.5.1320), KKIP3 / NufB4 (Tb927.10.6700), CPSF73 (Tb927.4.1340); a C-terminal 10× c-Ty to Pol-I RPA1 (Tb927.8.5090), RBP34 (Tb927.11.3340) and SNAP42 (Tb927.5.3910) and a C-terminal TurboID::4xGS::3xmyc to VEX1 (Tb927.11.16920) and VEX2 (Tb927.11.13380). pNAT^TAGx^ to add a N-terminal 6× c-myc to BDF2 (Tb927.10.7420), V2PL01 / ESB3 (Tb927.9.15850), V2PL04 / ESAP1 (Tb927.10.11600), KKIP2 / NufB4 (Tb927.5.1320), KKIP4 / NufB5 (Tb927.7.3080) and KKIP11 / NufB6 (Tb927.9.7820); and a N-terminal 3xmyc::4xGS::TurboID to VEX2 (Tb927.11.13380). TurboID [40] sequences were codon optimised for *T. brucei* expression [89, 90].

For the RNAi experiments, a previously generated plasmid was used to target VEX2 (Tb927.11.13380) [30]. For ESB1 (Tb927.10.3800), ESB2 (Tb927.1.1820), ESB3 (Tb927.9.15850), ESAP1 (Tb927.10.11600) RNAi constructs, primers were selected from ORF sequences using RNAit [91]. A specific RNAi target fragment was amplified and cloned in pRPa^iSL^ [88].

For tetracycline inducible ectopic expression, the ESB2 ORFs, either wild-type or containing an inactive nuclease (D240A, D330A and D353A), were cloned in the overexpression plasmid pRPa [93] containing a 12× c-myc tag at the C-terminus. A nuclear localisation (NLS) sequence (La; RGHKRSRE [92]) was added between the ORF and the 12× c-myc tag using HiFi cloning. Untagged versions of these plasmids were also generated. The ESB2 ORFs, either wild-type or containing an inactive nuclease, were recoded and synthesised (GenScript) to be refractory to the RNAi construct above described. To generate constructs for dual inducible overexpression and RNAi, the ESB2 RNAi stem-loop was then cloned into the above mentioned pRPa ESB2^recoded^ overexpression plasmids via Klenow.

The BES promoter-targeting construct, pESP::RFP::PAC, was derived from pESPi::RFP::PAC [93] by removing the tetracycline-operator and excised via PshAI / SacI prior to transfection.

RNAi, overexpression and dual RNAi / overexpression cassettes were excised via AscI prior to transfection. All plasmids were confirmed by whole plasmid sequencing using Plasmidsaurus.

### CRISPR/Cas9 precision editing

For CRISPR/Cas9 mediated precision editing, the sgRNAs were cloned into pT7^sgRNA^. The plasmid was sequenced for confirmation and then digested with NotI prior to transfection into a 2T1/T7/Cas9 parental cell line [86].

To edit either ESB2 D240 or D330, dsDNA templates ESB2 D240D, D240A, D240N, D330D, D330A and D330N were generated by end-filling reactions. Templates were identical except for the D240/D330 codons, which contained a synonymous mutation encoding aspartate, or mutations encoding alanine and asparagine, respectively. All templates contained 9-18 conservative changes in the regions immediately upstream and/or downstream of the target codons, flanked by 50 bp of complete homology to the wild-type sequence (a schematic representation is depicted in **Supplementary Fig. 8e**). All substitutions were based on preferred codon usage where possible [90].

Tetracycline was added to induce Cas9 induction, 48 hours later, ten million cells constitutively expressing a gRNA were transfected with ∼10-15 μg of repair template using Amaxa human T cell nucleofector solution (Lonza) and the programme Z-001. Twenty-four hours later, the cells were diluted to 1.5 cells/ml and plated in 96-well plates in the absence of any selection drug, incubated for 5 days, then 10-15 subclones for each mutation were expanded. To assess whether successful precision editing occurred, DNA was extracted with DNAzol and then analysed first by a diagnostic PCR using a reverse primer that specifically recognised the synonymous substitutions present in the ESB2 repair templates, therefore only generating an amplicon if the template had been integrated. For the positive clones, a PCR was performed to amplify a larger fragment using primers that bind regions outside the repair template and that should amplify fragments regardless of whether editing occurred - the resulting amplicons were purified and sent for Sanger sequencing to confirm the genotypes.

### TurboID-mediated proximity labelling

3xmyc::4xGS::TurboID::VEX2, VEX2::TurboID::4xGS::3xmyc and VEX1::TurboID::4xGS::3xmyc expressing parasites were grown to 5 × 10^5^ cell.ml^−1^, at which point biotinylation was initiated in four to six replicates of each cell line, by adding biotin to 50 μM for 18 h. The parental line treated with biotin and TurboID cell lines treated with DMSO (biotin vehicle) were used as controls. After *in vivo* biotinylation, parasites were harvested by centrifugation, washed three times in PBS/2% glucose and pellets were stored at −80 °C until lysis or processed right away.

The affinity purification of biotinylated material was adapted from [40, 94]. Briefly, a pellet of 5 × 10^8^ parasites was used for each affinity purification which was lysed in 1 mL of ice-cold RIPA buffer (50 mM Tris.HCl pH 7.4, 150 mM NaCl, 1% NP-40, 0.5% sodium deoxycholate, 0.1% SDS) containing 0.1 mM PMSF, 1.5 μM Pepstatin, 0.1 mM TLCK. Additionally, every 20 ml of RIPA was supplemented with 200 μl proteolytic protease inhibitor cocktail containing w/v 2.16% 4-(2-aminoethyl)benzenesulfonyl fluoride hydrochloride, 0.047% aprotinin, 0.156% bestatin, 0.049% E-64, 0.084% Leupeptin, 0.093% Pepstatin A (Abcam) and one tablet of complete protease inhibitor EDTA free (Roche). Lysates were sonicated using Bioruptor (settings high; 30 sec on / 30 sec off; 3 cycles) according to manufacturer’s instructions. One microlitre of microccocal nuclease was added to each lysate, digestion of nucleic acids proceeded for 10 mins at RT followed by 50 mins on ice. Lysates were clarified by centrifugation at 10,000 g for 10 min at 4 °C. For enrichment of biotinylated material, 100 μl of magnetic streptavidin bead suspension (1 mg of beads, Resyn Bioscience) was used for each affinity purification from 5 × 10^8^ parasites. Biotinylated material was affinity purified by end-over-end rotation at 4 °C overnight. Beads were washed in 1 mL of the following for 5 min each: RIPA for six washes, 4 M urea in 50 mM ammonium bicarbonate (AB) pH8.5 for two washes, 6 M urea in 50 mM AB pH8.5 for two washes, 1 M KCl in 50 mM AB pH8.5 for two washes and 50 mM AB pH8.5 for two washes.

4% of the material was used for protein blotting analysis to confirm successful enrichment for biotinylated material prior to proteomics analysis. The remaining 96% was processed as follows:

Beads from each affinity purification were then resuspended in 200 μl 50 mM TEAB pH8.5 containing 0.01% ProteaseMAX (Promega), 10 mM TCEP, 10 mM Iodoacetamide, 1 mM CaCl_2_ and 500 ng Trypsin Lys-C (Promega). The on-bead digest was carried out overnight at 37 °C while shaking at 200 rpm. Supernatant from digests was retained and beads were washed for 5 min in 50 μl water which was then added to the supernatant. Digests were acidified with trifluoroacetic acid (TFA) to a final concentration of 0.5% before centrifugation for 10 min at 17,000×*g*. The supernatant was desalted using C_18_ 0.6 μL ZipTips (Millipore), elution volume was 20 μl. Desalted peptides were dried for MS analysis.

### Mass spectrometry data acquisition

For total proteome TurboID analysis, peptides were loaded onto an mClass nanoflow UPLC system (Waters) equipped with a nanoEaze M/Z Symmetry 100 Å C_18_, 5 µm trap column (180 µm x 20 mm, Waters) and a PepMap, 2 µm, 100 Å, C_18_ EasyNano nanocapillary column (75 μm x 500 mm, Thermo). The trap wash solvent was aqueous 0.05% (v:v) trifluoroacetic acid and the trapping flow rate was 15 µL/min. The trap was washed for 5 min before switching flow to the capillary column. Separation used gradient elution of two solvents: solvent A, aqueous 0.1% (v:v) formic acid; solvent B, acetonitrile containing 0.1% (v:v) formic acid. The flow rate for the capillary column was 330 nL/min and the column temperature was 40°C. The linear multi-step gradient profile was either: 3-10% B over 7 mins, 10-35% B over 30 mins, 35-99% B over 5 mins and then proceeded to wash with 99% solvent B for 4 min; or 2.5-10% B over 10 mins, 10-35% B over 75 mins, 35-99% B over 15 mins and then proceeded to wash with 99% solvent B for 5 min. The same gradient was used for all samples that were directly compared. The column was returned to initial conditions and re-equilibrated for 15 min before subsequent injections.

The nanoLC system was interfaced with an Orbitrap Fusion hybrid mass spectrometer (Thermo) with an EasyNano ionisation source (Thermo). Positive ESI-MS and MS<u>2</u> spectra were acquired using Xcalibur software (version 4.0, Thermo). Instrument source settings were: ion spray voltage, 1,900 V; sweep gas, 0 Arb; ion transfer tube temperature, 275 °C. MS<u>1</u>spectra were acquired in the Orbitrap with 120,000 resolution, scan range: *m/z* 375– 1500; AGC target, 4e<u>5</u>; max fill time, 100 ms. Data-dependent acquisition was performed in top speed mode using a fixed 1 s cycle, selecting the most intense precursors with charge states 2–5. Easy-IC was used for internal calibration. Dynamic exclusion was performed for 50 s post precursor selection and a minimum threshold for fragmentation was set at 5e<u>3</u>. MS<u>2</u> spectra were acquired in the linear ion trap with: scan rate, turbo; quadrupole isolation, 1.6 *m/z*; activation type, HCD; activation energy: 32%; AGC target, 5e<u>3</u>; first mass, 110 *m/z*; max fill time, 100 ms. Acquisitions were arranged by Xcalibur to inject ions for all available parallelizable time.

### Mass spectrometry data analysis

Peak lists in.raw format were imported into Progenesis QI (Version 2.2., Waters) for peak picking and chromatographic alignment. A concatenated product ion peak list was exported in .mgf format for database searching against the *Trypanosoma brucei brucei* L427 subset of the TriTrypDB (11,388 sequences; 5,560,262 residues) database, appended with common proteomic contaminants. Mascot Daemon (version 2.6.1, Matrix Science) was used to submit searches to a locally-running copy of the Mascot programme (Matrix Science Ltd., version 2.7.0.1). Search criteria specified: Enzyme, trypsin; Max missed cleavages, 1; Fixed modifications, Carbamidomethyl (C); Variable modifications, Oxidation (M), Phospho (STY), Acetyl (Protein N-term), Biotin (Protein N-term, K); Peptide tolerance, 3 ppm (# 13 C = 1); MS/MS tolerance, 0.5 Da; Instrument, ESI-TRAP. Peptide identifications were passed through the percolator algorithm to achieve a 1% false discovery rate as assessed empirically by reverse database search, and individual matches were filtered to require minimum expected scores of 0.05. The Mascot.XML results file was imported into Progenesis QI, and peptide identifications associated with precursor peak areas were mapped between acquisitions. Relative protein abundances were calculated using precursor ion areas from non-conflicting unique peptides. For total proteome data, only non-modified peptides were used for protein-level quantification. Normalisation and statistical testing were performed in Progenesis QI, with the null hypothesis being peptides are of equal abundance among all samples [95].

Normalised protein label-free peak areas were exported from Progenesis LFQ and those proteins with at least two unique peptides identified were retained for downstream analysis. Missing values were imputed by drawing values from a left-shifted normal log2 intensity distribution to model low abundance proteins (VEX2 C-term, mean = 16.8, sd = 3.5; VEX2 N-term, mean = 16.8, sd = 3.5; VEX1, mean = 17.5, sd = 3.0). Proximal proteins were determined with the limma package [96] using options trend = TRUE and robust = TRUE for the eBayes function. Multiple testing correction was carried out according to Benjamini & Hochberg, the false discovery rate for identified proximals was 1%. GO term analysis was performed using TriTrypDB [97] and Revigo [98].

### Protein blotting

Protein samples were run according to standard protein separation procedures, using SDS-PAGE. 4-12% Bis-Tris gels were used (NuPAGE, Invitrogen). Otherwise, protein blotting was carried out according to standard protocols. The following primary antibodies were used: mouse α-myc (Millipore, clone 4A6, 1:10,000), mouse anti-Ty (BB2, Invitrogen, 1:5,000) and mouse α-EF1α (Millipore, clone CBP-KK1, 1:30,000). Streptavidin-HRP (Jackson ImmunoResearch) was used at 1:10,000 in PBS/5% BSA.

We used horseradish peroxidase coupled secondary antibodies (α-mouse and α-rabbit, Bio-Rad, 1:5,000). Blots were developed using an enhanced chemiluminescence kit (Amersham) according to the manufacturer’s instructions. Densitometry was performed using Fiji v. 2.9.0 [99].

### Immunofluorescence microscopy

Immunofluorescence microscopy was carried out according to standard protocols [33]. For wide-field microscopy, the cells were attached to 12-well 5 mm slides (Thermo Scientific). For super-resolution microscopy, the cells were attached to poly-L-lysine-treated high precision coverslips (thickness 1^1/2^ mm), stained and then mounted onto glass slides. Cells were mounted in Vectashield with DAPI (wide field) or stained with 1 µg mL^−1^ DAPI for 10 min and then mounted in Vectashield without DAPI (super-resolution). Primary antisera were rat α-VSG-2 (1:10,000), rabbit α-VSG-6 (1:10,000), rabbit α-myc (NEB, clone 71D10, 1:200), mouse α-myc (NEB, clone 9B11, 1:2,000), mouse α-Pol-I (largest subunit; 1:100 [34]) and mouse α-Ty (Invitrogen, BB2, 1:1,000). Streptavidin-Alexa fluor 488 was applied at 1:500 (Invitrogen).

The secondary antibodies were Alexa Fluor conjugated goat antibodies: α-mouse, α-rat and α-rabbit, Alexa Fluor 488, Alexa Fluor 555 Plus or Alexa Fluor 568 (1:1,000 for super-resolution microscopy or 1:2,000 for wide-field microscopy).

### Microscopy and image analysis

For wide-field microscopy, cells were analysed using a Zeiss AxioObserver Inverted Microscope equipped with Colibri 7 narrow-band LED system and white LED for epifluorescent and white light imaging and ZEN Pro software (Carl Zeiss). Images were acquired as *z*-stacks (0.1–0.2 μm) and further deconvolved using the default settings (‘good, medium’) in ZEN Pro. For super-resolution microscopy, cells were analysed using a Zeiss LSM980 Airyscan 2 or a Zeiss Elyra 7 and the Zeiss ZEN software (Carl Zeiss). Representative images obtained by super-resolution microscopy correspond to maximum 3D projections by the brightest intensity of stacks of approximately 30 slices of 0.1 μm. Images acquired with the Zeiss LSM980 Airyscan 2 were deconvolved using Airyscan Joint Deconvolution (XY resolution ∼90 nm). Super-resolution structured illumination microscopy (SR-SIM) was performed using a Zeiss Elyra 7 microscope in Lattice SIM^2^ mode (XY resolution ∼60 nm). SIM reconstruction was performed after correcting for chromatic aberrations using the channel alignment function in Zen and performing deconvolution using the default settings. 100 nm Tetraspeck beads (Thermo) were adhered to slides and were used to determine channel alignment for each experiment. DAPI-stained *T. brucei* nuclear and mitochondrial DNA were used as cytological markers for cell-cycle stage; one nucleus and one kinetoplast (1N:1K) indicates G1, one nucleus and an elongated kinetoplast (1N:eK) indicates S phase, one nucleus and two kinetoplasts (1N:2K) indicates G2/M and two nuclei and two kinetoplasts (2N:2K) indicates post-mitosis [100, 101]. All the images were processed and scored using Fiji v.2.9.0 [99]. Pearson’s correlation coefficient was applied as a statistical measure of colocalisation [102]. Overlapping, adjacent and separate foci presented a Pearson’s correlation coefficient in the ranges ≥0.5 to ≤1, ≥−0.5 to <0.5 and ≥−1 to <−0.5, respectively. Counts in total cells or specific cell cycle phases were typically performed using >100 nuclei. All quantifications are averages or representative of at least two biological replicates and independent experiments.

### Transcriptomics

RNA-seq analysis was performed in bloodstream forms using 2T1 cells and uninduced or induced clones of ESB2 RNAi (24 and 36 h), ESB3 RNAi (24 and 36 h), ESAP1 RNAi (24 and 36 h), ESB2 wild-type overexpression (72 h) and ESB2 ‘inactive nuclease’ overexpression (72 h) – at least 3 biological replicates each. RNA-seq analysis was also performed in procyclic forms using PT1 cells and uninduced or induced clones of ESB2 RNAi (72 h), and ESAP1 RNAi (72 h) - 2 biological replicates each. RNA was extracted using the RNeasy kit (Qiagen) according to the manufacturer’s instructions. The RNA samples were sent for sequencing to BGI (Hong Kong). Briefly, polyadenylated transcripts were enriched using poly-dT beads and reverse-transcribed before sequencing on a DNBSeq platform. Each sample generated approximately 40 million reads (paired end, 100 bp) or 100 million reads (paired end, 150 bp).

Reads were mapped to either *T. brucei* L427 2018 [8] or a hybrid assembly consisting of the *T. brucei* 927 reference genome [103] plus the bloodstream *VSG*-ESs [9] and metacyclic *VSG-*ESs [104, 105] from the Lister 427 strain. In the figures, we show the data mapped to the hybrid assembly because individual *ESAG* annotations are more amenable. The gene cohorts affected by the knockdowns or overexpression experiments we performed are the same and affected to the same extent irrespective of which assembly was used for the analysis. Bowtie 2-mapping [106] was with the parameters -- very-sensitive --no-discordant --phred33. Alignment files were manipulated and filtered for MapQ1 or MapQ10 in PCFs and BSFs, respectively with SAMtools [107] - indeed, high stringency mapping was applied to distinguish *ESAG* transcripts. Bam files were inspected on Artemis [108] or IGV [109]. Per-gene read counts were derived using subread [110]. RPKM values were derived from normalised read counts and differential expression analysis was conducted with edgeR [111]. Coverage maps were generated using deepTools [112].

### RT-qPCR

RT-qPCR analysis was performed to determine *RFP, ESB2* and *ESAG7* mRNA levels in 2T1 bloodstream form cells and uninduced or induced clones of ESB2 RNAi (24 h) with a pESP::RFP::PAC reporter at the active-*VSG* expression-site – three biological replicates were analysed with three technical replicates per condition.

RT-qPCR analysis was also performed to determine *ESB2* mRNA levels in 2T1 bloodstream form cells and uninduced or induced clones of ESB2 RNAi (24 h), overexpression of either wild-type or containing an inactive nuclease (48, 72 h) or dual RNAi + overexpression (48 h), – two or three biological replicates were analysed with three technical replicates per condition.

RNA was extracted using the RNeasy kit (Qiagen) according to the manufacturer’s instructions. Reverse transcription of RNA and quantitative PCR was performed using the Luna Universal One-step RT-qPCR Kit using 5 ng or 25 ng of RNA per reaction, for *RFP/ESAG7* or *ESB2* related amplifications, respectively. Reactions were carried out in a QuantStudio 3 or QuantStudio 7 real time PCR system (ThermoFisher) and typically included reverse transcription at 55°C for 10 minutes, followed by initial denaturation at 95°C for 1 minute followed by 40 cycles of 95°C for 10 seconds and 60°C for 30 seconds with a signal read at the end of each cycle plus a final melting curve to check fidelity from 60-95°C, with a signal read every 1°C. The data was normalised against the *T. brucei* actin gene. Fold changes in mRNA abundance were calculated using the ΔΔCT method, normalised to the parental strain *T. brucei* 2T1 and displayed as relative abundance.

### Phylogenetic analysis and structural prediction

DNA and amino acid sequences of ESB2, ESB3 and ESAP1 and respective kinetoplastid orthologues were obtained from the TriTryp database [97] The remaining protein sequences were obtained from Uniprot [113]. Phylogenetic analysis was performed using TreeViewer [114]. Structural predictions were performed with AlphaFold2 optimised for trypanosomatid proteins [115, 116]. Sequence alignments were generated with Clustal W [117] and displayed in ESPript [118] or UCSF ChimeraX [119]. Protein structures were visualised and superimposed in UCSF ChimeraX. Sequence motif analysis was performed using InterPro [120] and structure-based domain analysis was performed using FoldSeek [121].

### Data visualisation & Statistics

Heatmaps, radial, volcano, scatter and violin plots were generated in RStudio using the following packages: ggplot2, RColorBrewer, ggrepel, dplyr, scales and tidyverse [122]. Growth curves, heatmaps, box plots, floating bar and stacked bar graphs were generated using GraphPad Prism Software (version 10.0). The GO term network analysis was generated using Cytoscape [123]. All statistical analyses were performed using GraphPad Prism Software (version 10.0), except for the transcriptomic and proteomics analyses, which were conducted using R (details provided in the respective sections).

### Resources and reagents

All unique materials are available on request.

## Data and code availability

RNA-Seq data was deposited in ENA (PRJEB89423). Proteomics data was deposited in ProteomeXchange (PXD063534) and can be downloaded from MassIVE (MSV000097776).

## Acknowledgements

This work was supported by a Wellcome Trust/Royal Society Sir Henry Dale Fellowship to JFRC (222573/Z/21/Z). We would like to thank the Imaging & Cytometry, Genomics and Proteomics & Metabolomics Labs in the Bioscience Technology Facility at the University of York. The York Centre of Excellence in Mass Spectrometry was created thanks to a major capital investment through Science City York, supported by Yorkshire Forward with funds from the Northern Way Initiative, and subsequent support from EPSRC (EP/K039660/1; EP/M028127/1). We thank Dr Vincent Geoghegan for advice regarding the proximity labelling experiments and Beth Spink for assistance in cell line maintenance. We thank Dr Sebastian Hutchinson and Prof David Horn for kindly gifting the PT1 cell line. Further, we would also like to acknowledge Prof Keith Gull, FRS, and all members of the Mottram and Cayla labs for helpful discussions.

## Contributions

LIML, JRCF conceived the study. LIML, SL, MJ, JRCF planned and performed experiments. LIML, LW, SL, MJ, AD, JRCF analysed data. JRCF supervised the study and secured funding. JRCF wrote the manuscript with contributions from LIML, LW and AD.

## Ethics declarations

The authors declare no competing interests.

---

**Supplementary Fig. 1.**
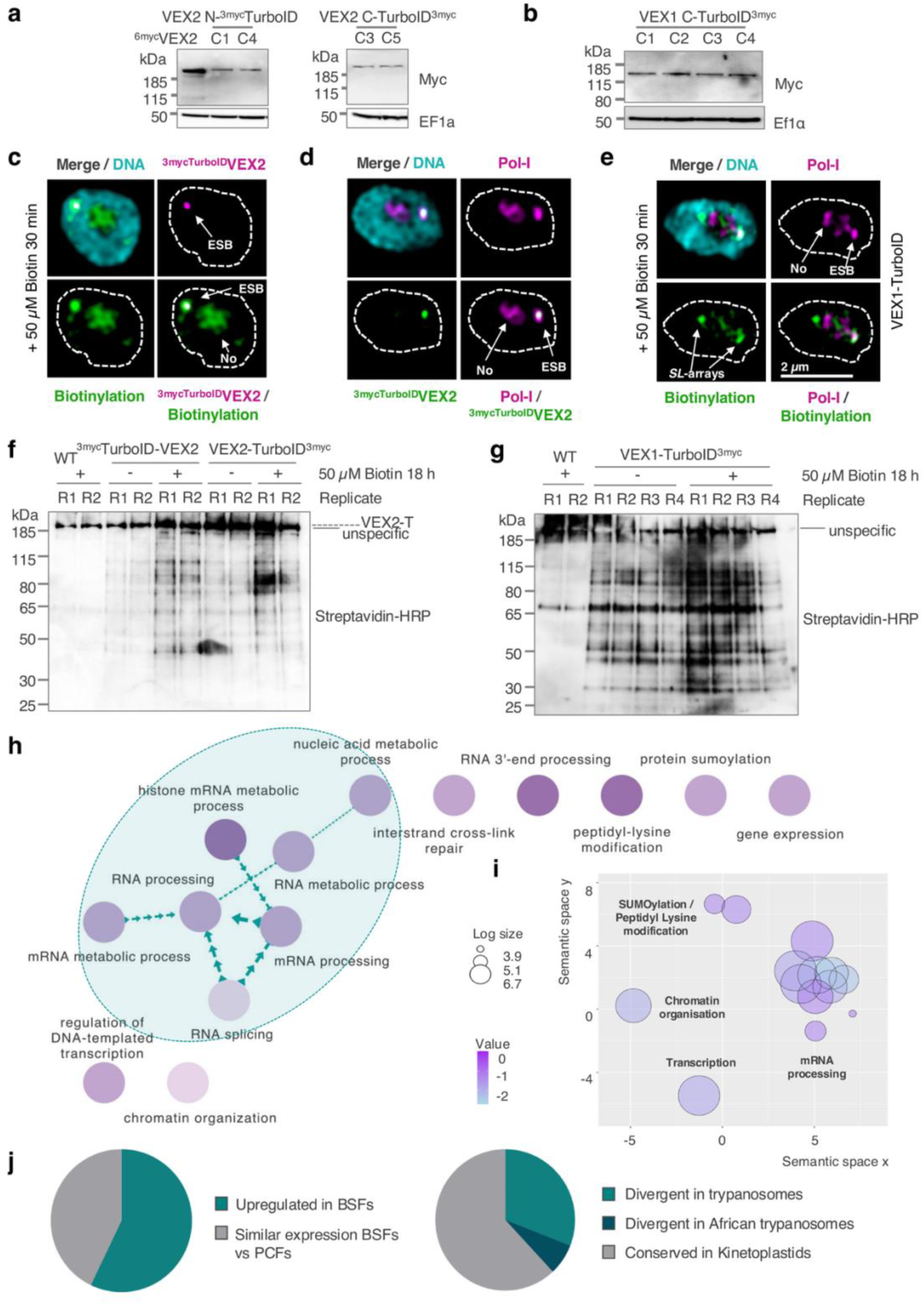
Characterisation of VEX-TurboID cell lines and ‘hit’ prioritisation for subsequent validation. **a-b**, protein blotting analysis of whole cell extracts of ^3myc^TurboID-VEX2 and VEX2-TurboID^3myc^ (**a**) and VEX1-TurboID^3myc^ (**b**) cell lines. An anti-myc antibody was used; EF1α, loading-control. *C* means clone x. **c-e,** fluorescence microscopy analysis of ^3myc^TurboID-VEX2 / Biotinylation (**c**), ^3myc^TurboID-VEX2 / Pol-I (**d**) and Biotinylation / Pol-I in a VEX1-TurboID^3myc^ cell line (**e**). Biotinylated material was detected using Streptavidin-A488. Images were acquired using a Zeiss LSM980 Airyscan 2 and correspond to 3D projections by brightest intensity of 0.1 μm stacks. DNA was stained with DAPI (cyan); scale bars: 2 μm. **f-g,** protein blotting analysis of immunoprecipitation samples of biotinylated material using streptavidin beads from ^3myc^TurboID-VEX2 and VEX2-TurboID^3myc^ (**f**) and VEX1-TurboID^3myc^ (**g**) cell lines treated or untreated with biotin. Streptavidin-HRP was used for detection. **h-j,** GO terms (**h, i**), protein expression and sequence homology (**j**) analyses of the 49 proteins significantly enriched in the VEX-TurboID ‘plus biotin’ samples that have not been previously associated with the ESB or surrounding nuclear bodies. **h**, protein network analysis based on biological process. **i**, similar GO terms containing functionally-related proteins cluster proximally. The size and colour of the bubbles were derived from the number of proteins in the VEX interactome that cluster in each GO term and the corresponding *p* value, respectively.

**Supplementary Fig. 2.**
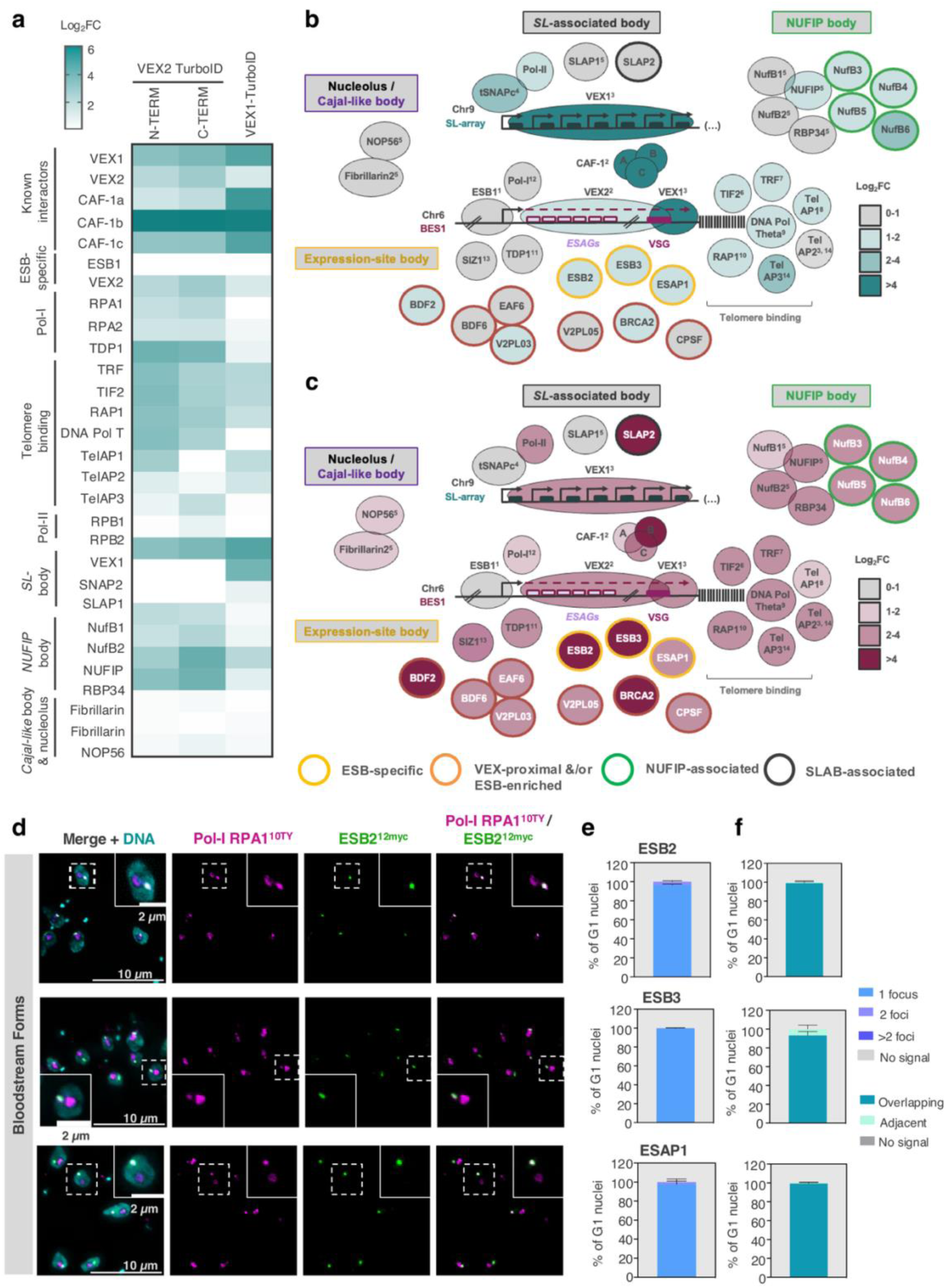
VEX1 and VEX2 TurboID captured most known components of the ESB and spatially proximal bodies and identified new components. **a**, heatmap depicting the Log_2_FC for ESB, SLAB, Cajal-body and NUFIP body associated factors in the ^3myc^TurboID-VEX2, VEX2-TurboID^3myc^ and VEX1-TurboID^3myc^ datasets. Log_2_FC represents the fold change in protein abundance between the cell lines where VEX1/VEX2 were fused with TurboID and the parental line, both treated with 50 μM of biotin for 18 hours. **b-c,** cartoon depicting the assembly of nuclear bodies that sustain *VSG* expression in *T. brucei* bloodstream forms. All known and newly identified factors are depicted and coloured according to their Log_2_FC in the VEX1-TurboID^3myc^ (**b**) and ^3myc^TurboID-VEX2 / VEX2-TurboID^3myc^ (**c**) datasets. Newly identified factors are outlined in yellow, orange, green or dark grey depending on their localisation. VEX1, VEX2 and ESB1 positions on the DNA are based on available ChIP-Seq data (Faria et al, 2019 & 2023; Lopez-Escobar et al, 2022). BDF6, EAF6 and V2PL03 (Tb927.10.14190) are known to interact (Staneva et al, 2021). For the molecules previously associated with the ESB, or surrounding bodies, the superscript numbers indicate the corresponding reference as follows: 1. López-Escobar et al., 2022, PMID: 35879525. 2. Faria et al., 2019, PMID: 31289266. 3. Glover et al., 2016, PMID: 27226299. 4. Das et al., 2005, PMID: 16055739. 5. Budzak et al., 2022, PMID: 35013170. 6. Jehi et al., 2014, PMID: 24810301. 7. Jehi et al., 2016, PMID: 27258069. 8. Reis et al., 2018, PMID: 29385523. 9. Leal et al., 2020, PMID: 32890403. 10. Yang et al., 2009, PMID: 19345190. 11. Narayanan and Rudenko, 2013, PMID: 23361461. 12. Navarro and Gull, 2001, PMID: 11742402. 13. López-Farfán, Bart et al. 2014, PMID: 25474309. 14. Weisert et al, 2024, PMID: 39681615. **d-f,** fluorescence microscopy analysis of ESB2^12myc^, ^6myc^ESB3, ^6myc^ESAP1 and Pol-I RPA1^10ty^ in bloodstream forms. Images were acquired using a Zeiss AxioObserver and correspond to 3D projections by brightest intensity of 0.1 μm stacks. DNA was stained with DAPI (cyan); scale bars: 2 or 10 μm. **e/f** show the % of G1 nuclei with 1, 2 or >2 ESB2/ESB3/ESAP1 foci (**e**) and % of G1 nuclei where ESB2/ESB3/ESAP1 are overlapping, immediately adjacent or separate from the ESB (**f**). Error bars indicate standard deviation. For more details, see section ‘Microscopy and image analysis’. The graphs depict averages of two biological replicates; >100 G1 cells per condition were analysed.

**Supplementary Fig. 3.**
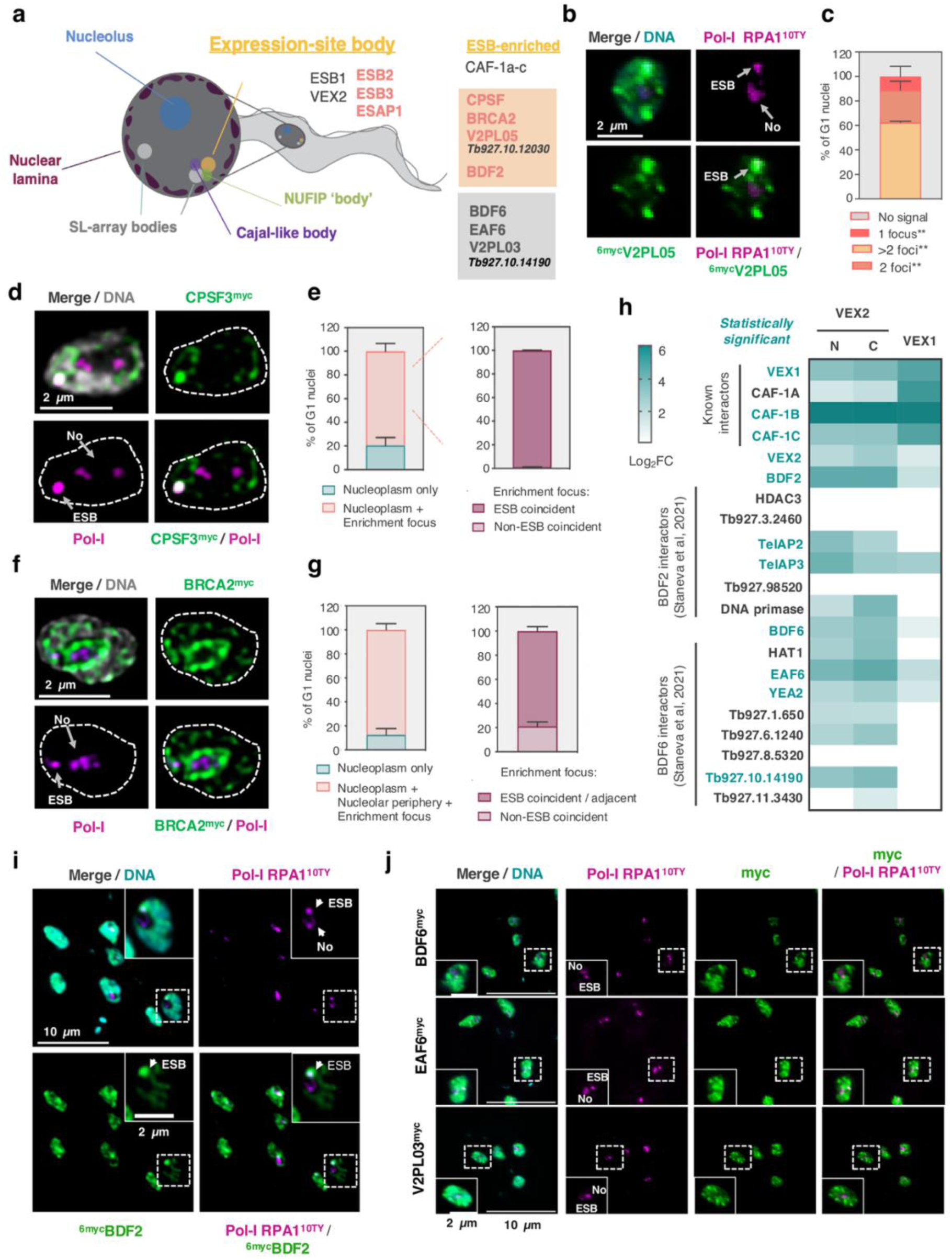
VEX proximity labelling identified nuclear factors that are significantly enriched at the ESB, including Tb927.10.12030, CPSF, BRCA2 and BDF2. **a**, cartoon summarising the main nuclear compartments in *T. brucei* bloodstream forms and highlighting the newly identified ESB-specific or ESB-enriched factors in salmon. BDF6, EAF6 and V2PL03, despite being found in proximity to VEX-proteins, did not evidently accumulate at the ESB. Created with BioRender.com **b-g / i-j,** fluorescence microscopy analysis of ^6myc^V2PL05 (Tb927.10.12030) (**b/c**), CPSF3^12myc^ (**d/e**), BRCA2^12myc^ (**f/g**), ^6myc^BDF2 (**i**), BDF6^12myc^, EAF6^12myc^ and V2PL03^12myc^ (Tb927.10.14190) (**j**) and Pol-I RPA1^10ty^ in bloodstream forms. **c** shows the % of G1 nuclei with 1, 2 or >2 ^6myc^V2PL05 foci. The asterisk indicates that additional nucleoplasmic signal can be detected. **e/g** show % of G1 nuclei where a major CPSF3 or BRCA2 enrichment focus is ESB coincident or not. For more details, see section ‘Microscopy and image analysis’. **c/e/g,** the graphs depict averages of two biological replicates; >100 G1 cells per condition were analysed; error bars indicate standard deviation. The images in **b/i/j** were acquired using a Zeiss AxioObserver, whereas the images in **d/f** were acquired using a Zeiss LSM980 Airyscan 2 and a Zeiss Elyra 7, respectively. All correspond to 3D projections by brightest intensity of 0.1 μm stacks. DNA was stained with DAPI (cyan or grey); scale bars: 2 μm or 10 μm (as indicated). ESB, expression-site body; No, nucleolus. **h,** heatmap depicting the Log_2_FC for BDF2 and BDF6 interactors in the ^3myc^TurboID-VEX2, VEX2-TurboID^3myc^ and VEX1-TurboID^3myc^ datasets. Log_2_FC represents the fold change in protein abundance between the cell lines where VEX1/VEX2 were fused with TurboID and the parental line, both treated with 50 μM of biotin for 18 hours.

**Supplementary Fig. 4.**
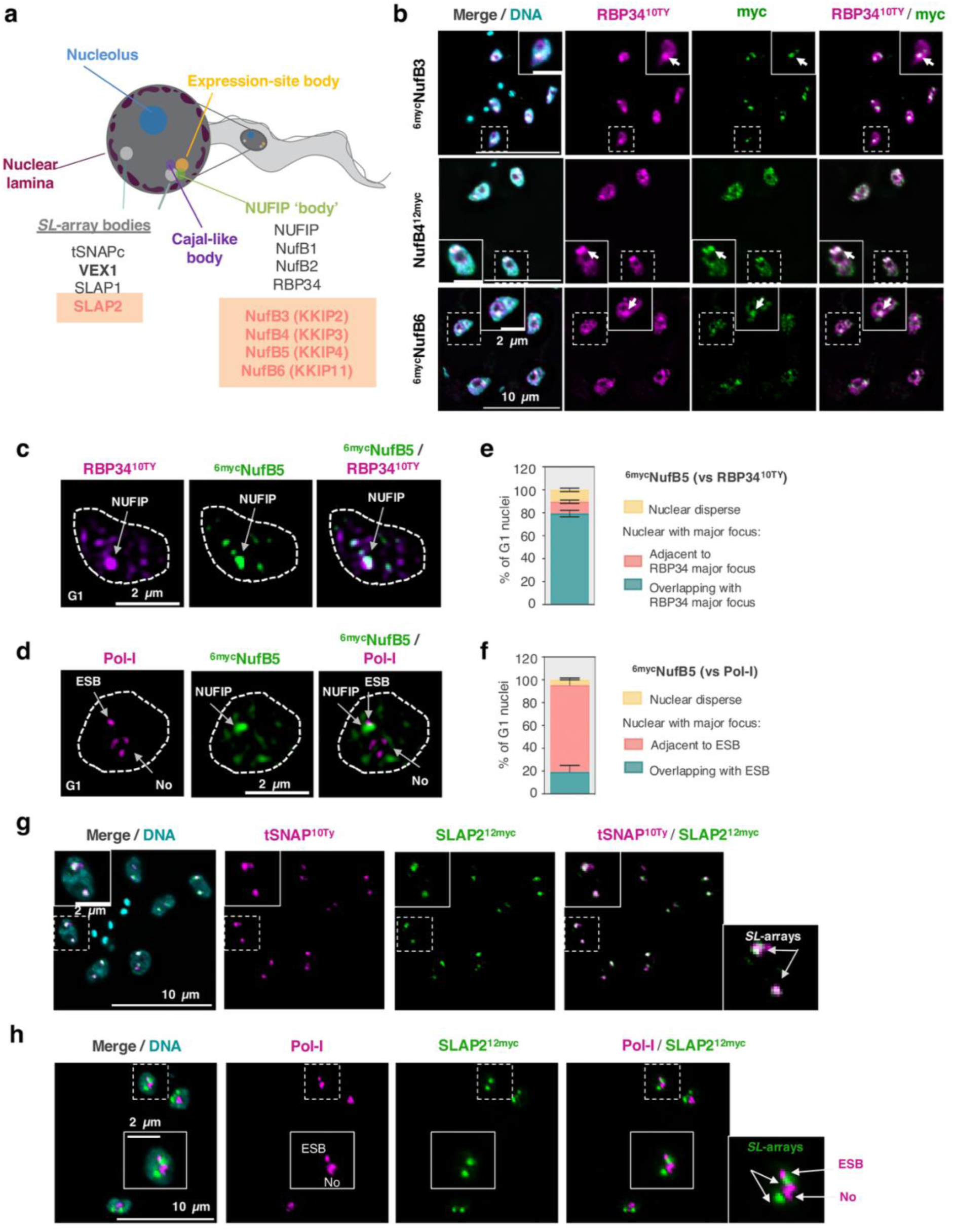
VEX proximity labelling identified new SLAB and NUFIP body components. **a**, cartoon summarising the main nuclear compartments in *T. brucei* bloodstream forms and highlighting the newly identified SLAB and NUFIP-body components in salmon. Created with BioRender.com **b-f,** fluorescence microscopy analysis of ^6myc^NufB3 (**b**), NufB4^12myc^ (**b**), ^6myc^NufB5 (**c/d**), ^6myc^NufB6 (**b**) and Pol-I RPA1^10ty^ or RBP34^10ty^ in bloodstream forms. **e/f,** the graphs depict averages of two biological replicates; >100 G1 cells per condition were analysed; error bars indicate standard deviation. For more details, see section ‘Microscopy and image analysis’. **g-h,** fluorescence microscopy analysis of SLAP2^12myc^ and tSNAP^10ty^ (**g**) or Pol-I RPA1^10ty^ (**h**) in bloodstream forms. The images in **b/g/h** were acquired using a Zeiss AxioObserver, whereas the images in **c/d** were acquired using a Zeiss LSM980 Airyscan 2. All correspond to 3D projections by brightest intensity of 0.1 μm stacks. DNA was stained with DAPI (cyan or grey); scale bars: 2 μm or 10 μm (as indicated). ESB, expression-site body; No, nucleolus.

**Supplementary Fig. 5.**
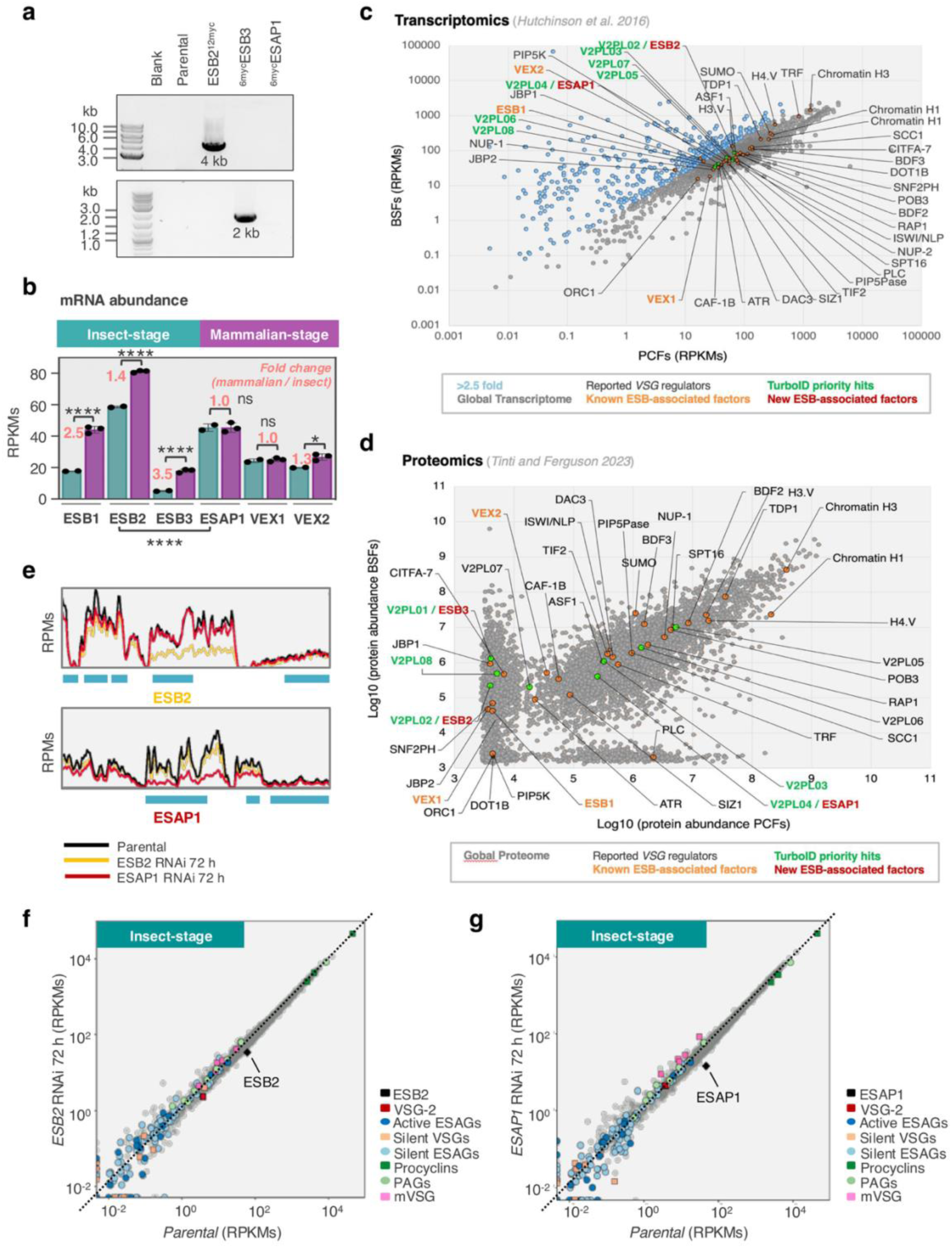
ESB2, ESB3 and ESAP1 are developmentally regulated. **a**, PCR analysis of the integration of ESB2 and ESB3 tagging constructs in the correct genomic location in procyclic forms. **b,** mRNA abundance expressed in reads per kilobase per million (RPKMs) in bloodstream (n=3, purple) and procyclic (n=2, cyan) forms. Data was obtained from RNA-Seq analysis of both developmental stages of the parasite. Statistical significance was determined using One-way ANOVA; ns, non-significant, **** *p*<0.0001. Error bars indicate standard deviation. **c-d,** Scatter plots represent either mRNA abundance (**c**) or protein abundance (**d**) in procyclic forms (x axis) and bloodstream forms (y axis). RNA-Seq data Hutchinson et al, 2016; proteomics data from Tinti & Ferguson, 2023. Labelled in grey are all the factors that have somehow been implicated in *VSG* regulation; in orange those that had previously been shown to localise to the ESB and in red the new ones identified in this study. GeneIDs for all TurboID priority hits can be found in the methods section. **e-g,** RNA-Seq analysis of ESB2 and ESAP1 knockdown in procyclic forms at 72 hours post induction. Reads per kilobase per million (RPKMs) are averages of two biological replicates. **e** zooms in on the ESB2 and ESAP1 coding loci to show successful transcript depletion in the knockdowns. RPMs, reads per million.

**Supplementary Fig.6.**
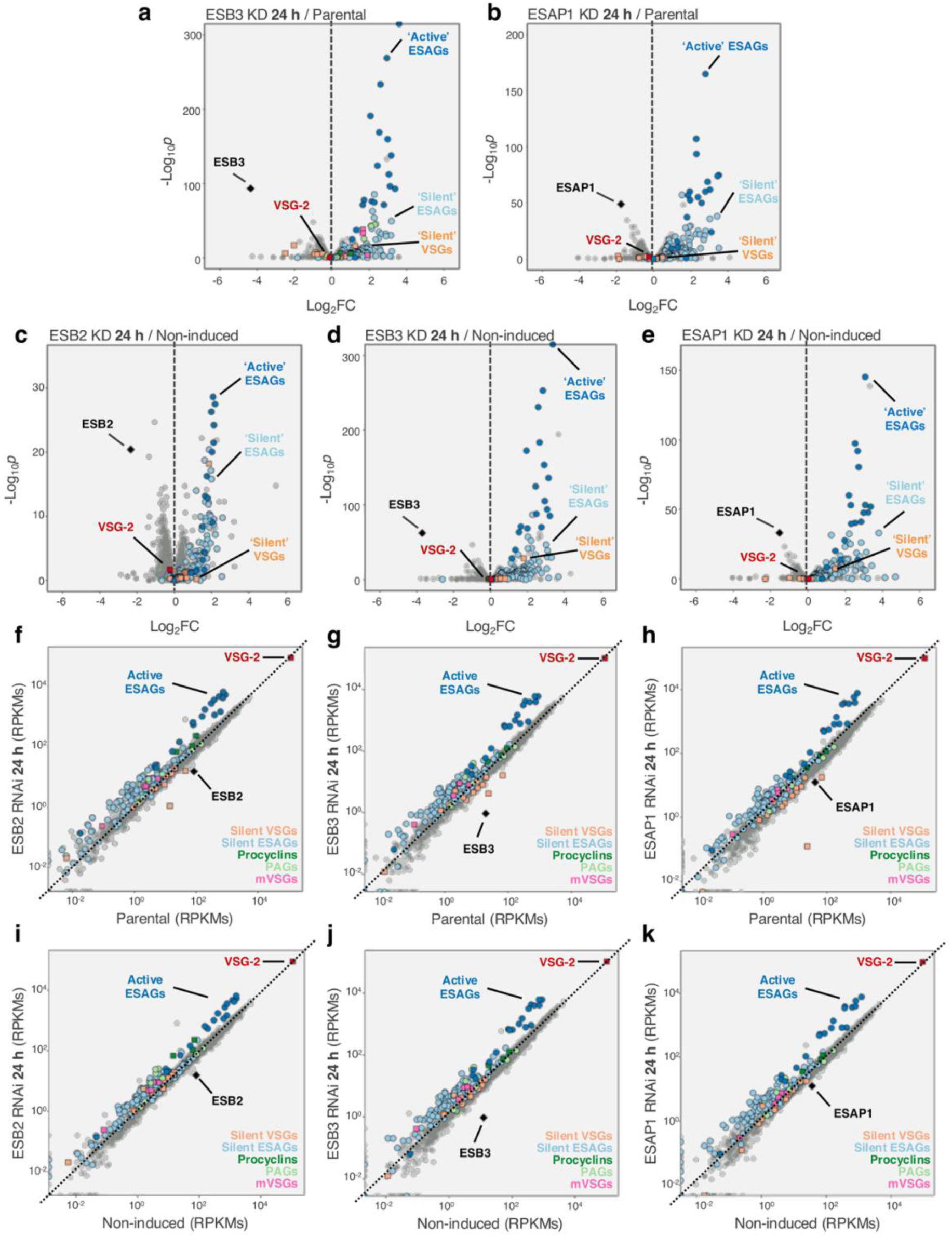
RNA-Seq analysis of ESB2, ESB3 and ESAP1 RNAi in BSFs. **a-k**,RNA-Seq analysis of ESB2 (n=5), ESB3 (n=3) and ESAP1 (n=3) knockdowns in bloodstream forms (BSFs) at 24 hours post induction, where *n* values correspond to the number of biological replicates. **a-e,** volcano plots depicting Log_2_FC (fold change) between RNAi and parental line (**a-b**) or RNAi induced *versu*s non-induced samples (**c-e**) and corresponding statistical significance (-log_10_*p*). The following cohorts are highlighted: ‘active’ *VSG* (red) and *ESAGs* (darker blue) and ‘silent’ *VSGs* (orange) and *ESAGs* (lighter blue). **f-k,** scatter plots comparing induced RNAi samples with the parental line (**f-h**) or the non-induced samples (**i-k**). RPKMs, reads per kilobase per million. The following cohorts are highlighted: ‘active’ *VSG* (red) and *ESAGs* (darker blue); ‘silent’ *VSGs* (orange) and *ESAGs* (lighter blue), metacyclic *VSGs* (mVSGs, pink), procyclins (darker green), procyclin associated genes (PAGs, lighter green).

**Supplementary Fig. 7.**
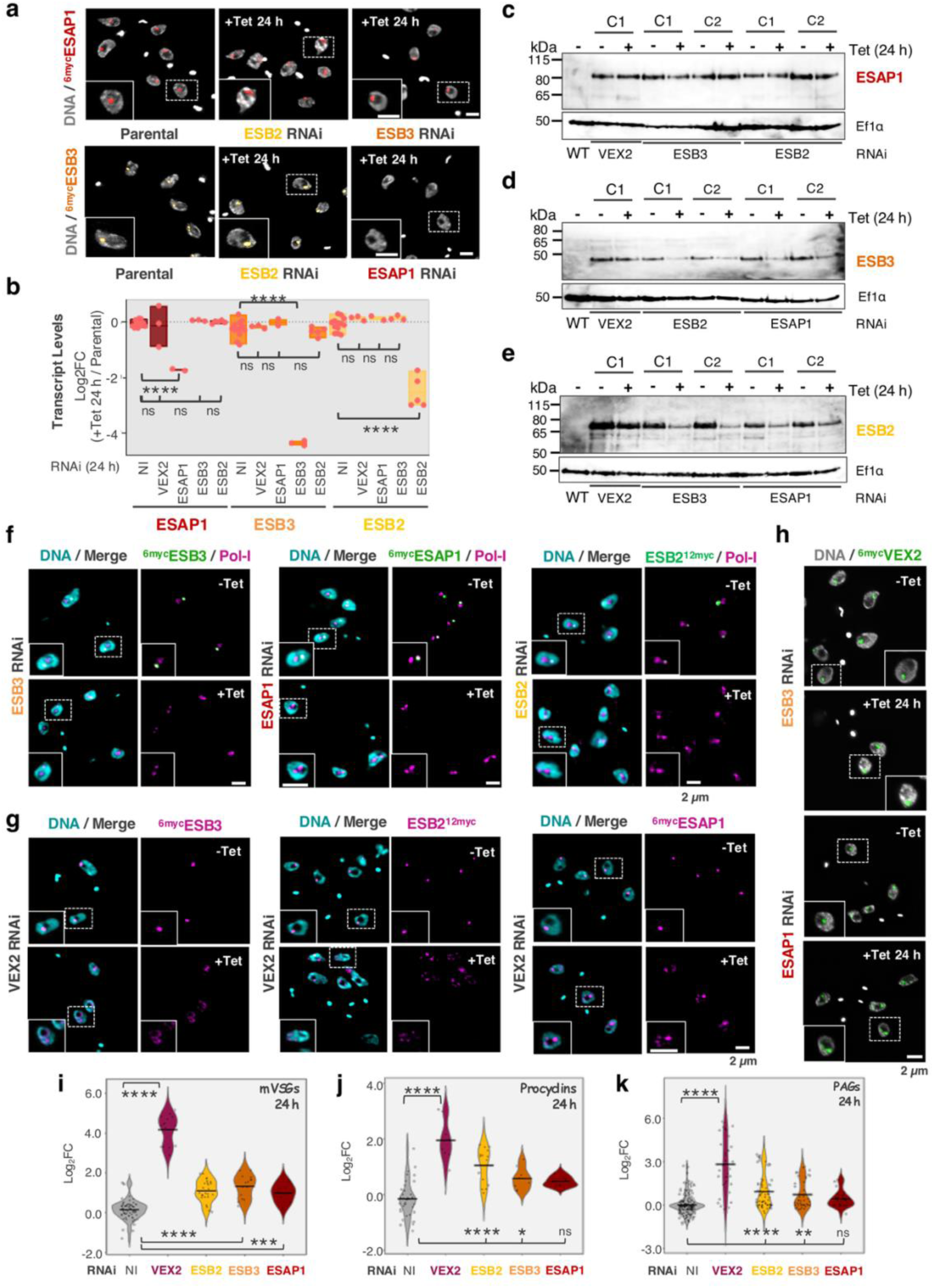
ESB2, ESB3, ESAP1, VEX2 and Pol-I co-dependencies regarding subcellular localisation as well as mRNA / protein abundance. **a**, Fluorescence microscopy analysis of ^6myc^ESB3 localisation following ESB2 and ESAP1 RNAi and ^6myc^ESAP1 localisation following ESB2 and ESB3 RNAi. **b,** ESB2, ESB3 and ESAP1 mRNA levels following VEX2, ESB2, ESB3 and ESAP1 RNAi. The values were derived from RNA-Seq data (ESB3 & ESAP1 RNAi: 3 biological replicates; ESB2 RNAi: 5 biological replicates). Bars span between minimum and maximum values; the line corresponds to the mean value, all datapoints are represented. Two-tailed paired Student’s *t* tests were applied; ns, non-significant; \*\*\*\**p* < 0.0001. **c-e,** protein blotting analysis of ^6myc^ESAP1 (**c**), ^6myc^ESB3 (**d**) and ESB2^12myc^ (**e**) levels following VEX2, ESB2, ESB3 or ESAP1 RNAi at 24 hours post induction. C1, clone 1; C2, clone 2. EF1α, loading-control. **f-h,** fluorescence microscopy analysis of Pol-I localisation following ESB2, ESB3 and ESAP1 knockdown (**f**), ESB2^12myc^, ^6myc^ESB3 and ^6myc^ESAP1 localisation following VEX2 knockdown (**g**) and ^6myc^VEX2 localisation following ESB3 and ESAP1 knockdown (**h**). **a/f-h,** all analyses were performed at 24 hours post induction. The images were acquired using a Zeiss AxioObserver and correspond to 3D projections by brightest intensity of 0.1 μm stacks. DNA was stained with DAPI (grey or cyan); scale bars: 2 μm. **i-k**, violin plots depicting a comparative transcriptomic analysis between VEX2, ESB2, ESB3 and ESAP1 RNAi cell lines with a focus on Pol-I transcribed gene cohorts: metacyclic *VSGs* (**d**), procyclins (**e**) and procyclin associated genes (*PAGs)* (**f**). Log_2_FC, fold change in transcript abundance between RNAi (24 hours post induction) and the parental cell line. The violins span between minimum and maximum values, centre lines correspond to the mean, all datapoints are shown. One-way ANOVA; ns, non-significant; * *p*<0.05; ** *p*<0.01; *** *p*<0.001; **** *p*<0.0001.

**Supplementary Fig. 8.**
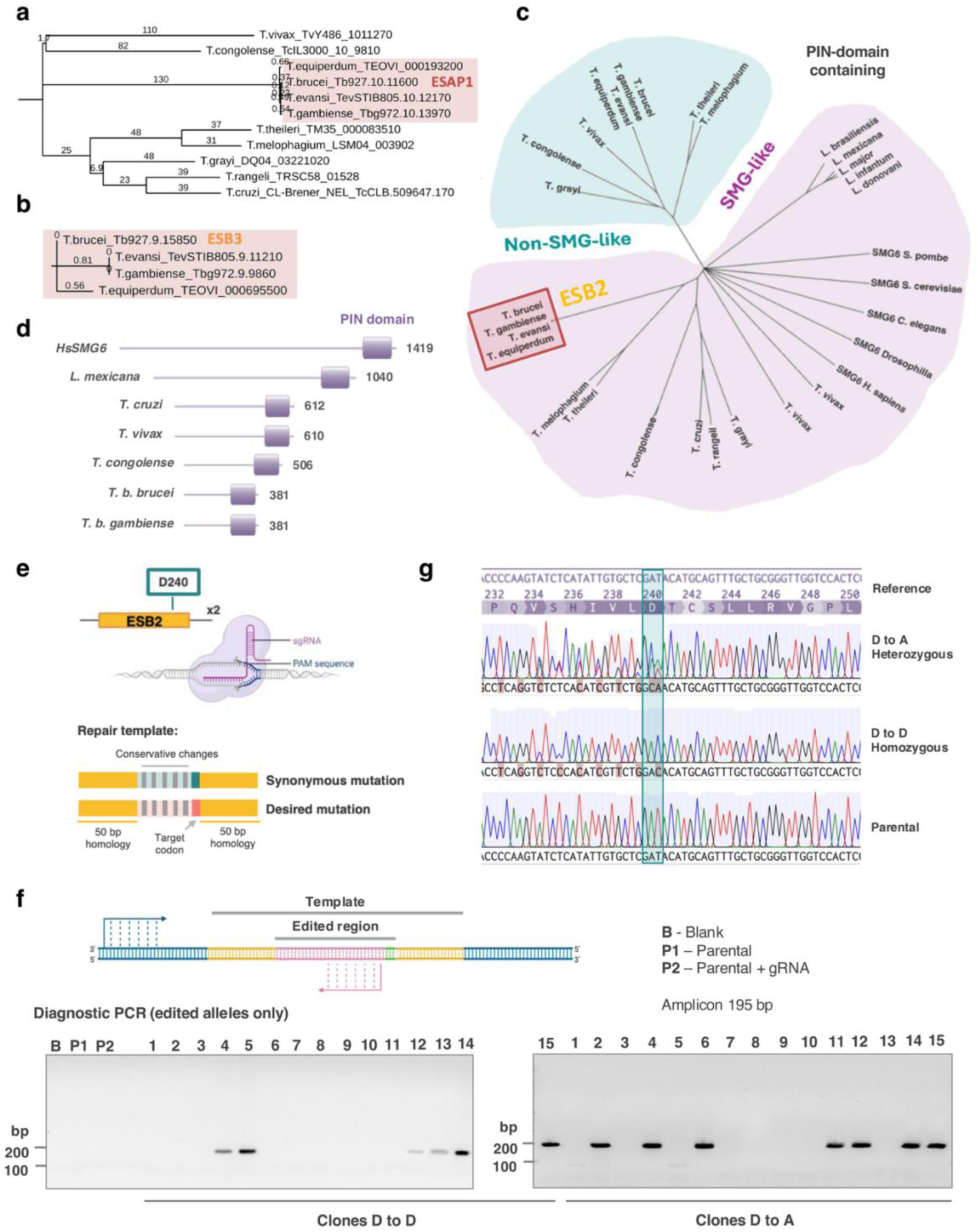
ESB2, ESB3 and ESAP1 phylogenetic analysis and ESB2 D240 precision editing. **a-c**, phylogenetic analysis of ESAP1 (**a**), ESB3 (**b**) and ESB2 (**c**) conducted using TreeViewer. Highlighted in light red are trypanosome species that contain *ESAGs*. **d,** cartoon depicting the domain architecture of SMG-like PIN containing proteins in trypanosomatids compared to human SMG6. **e-g,** CRISPR/Cas9 mediated precision editing of ESB2 D240. Mutants carrying the mutation from aspartic acid to alanine were always heterozygous, however successful double allele editing was achieved when using a repair template containing a synonymous mutation. Both repair templates contained conservative changes so that edited alleles could be readily distinguished from wild-type alleles (**e**). 15 clones were screened for both synonymous and non-synonymous mutations, respectively. **f,** diagnostic PCR analysis. The Sanger sequencing profiles depicted in **g** are representative of different clones and two independent experiments.

**Supplementary Fig. 9.**
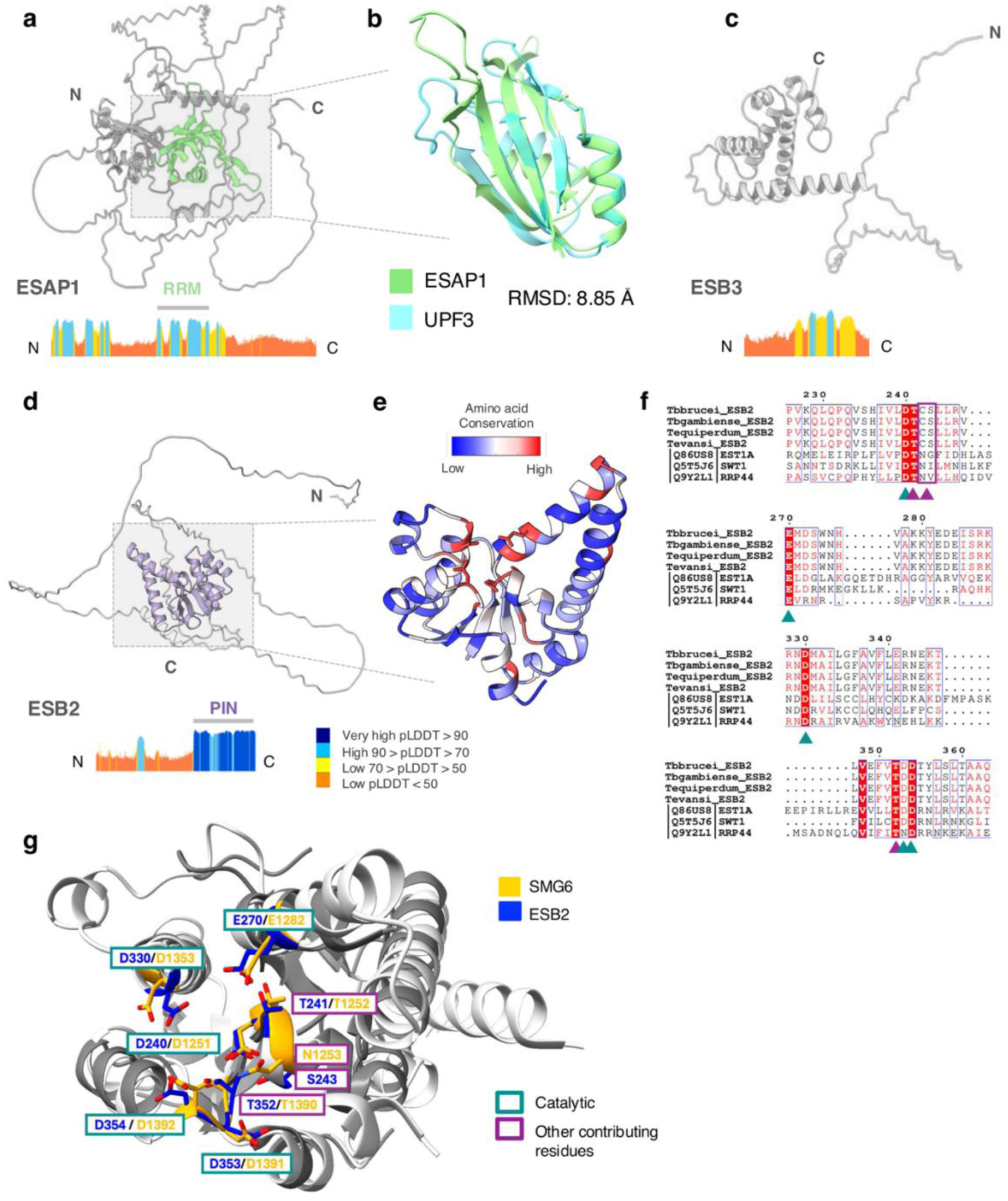
ESB2 and ESAP1 share structural features with nonsense mediated decay (NMD) factors. **a/c/d** depict 3D structural predictions of ESAP1 (RRM-like domain highlighted in green), ESB3 and ESB2 (PIN domain highlighted in purple). pLDDT plots accompany each model. **b,** Structural overlay between ESAP1 and UPF3 (PDB ID 7NWU). Root mean square deviation (RMSD) quantifies the average distance between corresponding atoms in two superimposed structures. **e** shows a structural prediction of ESB2 PIN domain overlayed with the structure of SMG6 PIN domain (PDB ID 2HWW) coloured by degree of amino acid conservation. **f-g,** conservation of the catalytic (cyan) and other important residues (purple) within the PIN domain of ESB2 compared to human SMG6. The first conserved, acidic residue, which is invariably an aspartate occurs at the end of β1 before the first helix (D240 / D1251). This is followed by another important residue, which always occurs after one turn of the first helix (α1) and often is an asparagine or a serine (Ser243 / N1253). The third residue of the active site (2^nd^ acidic residue) occurs in the helix (α2) following β2 and is always a glutamate (E270 / E1282); structurally is the most distant of the core acidic residues. Its side chain is held in place by the residue located between the first two active site residues, which is often a threonine (T241 / T1252). The fourth residue (3^rd^ acidic residue) is always an aspartate (D330 / D1353) and it is located on the other side of the active site at the end of α3. Finally, the active site contains one or sometimes two additional aspartic acid residues, which both occur in the loop following β4 and in the beginning following helix (D353 / D1391; D354 / D1392). Another potentially important residue in this region is often either a serine or threonine (T352 / T1390).

**Supplementary Fig. 10.**
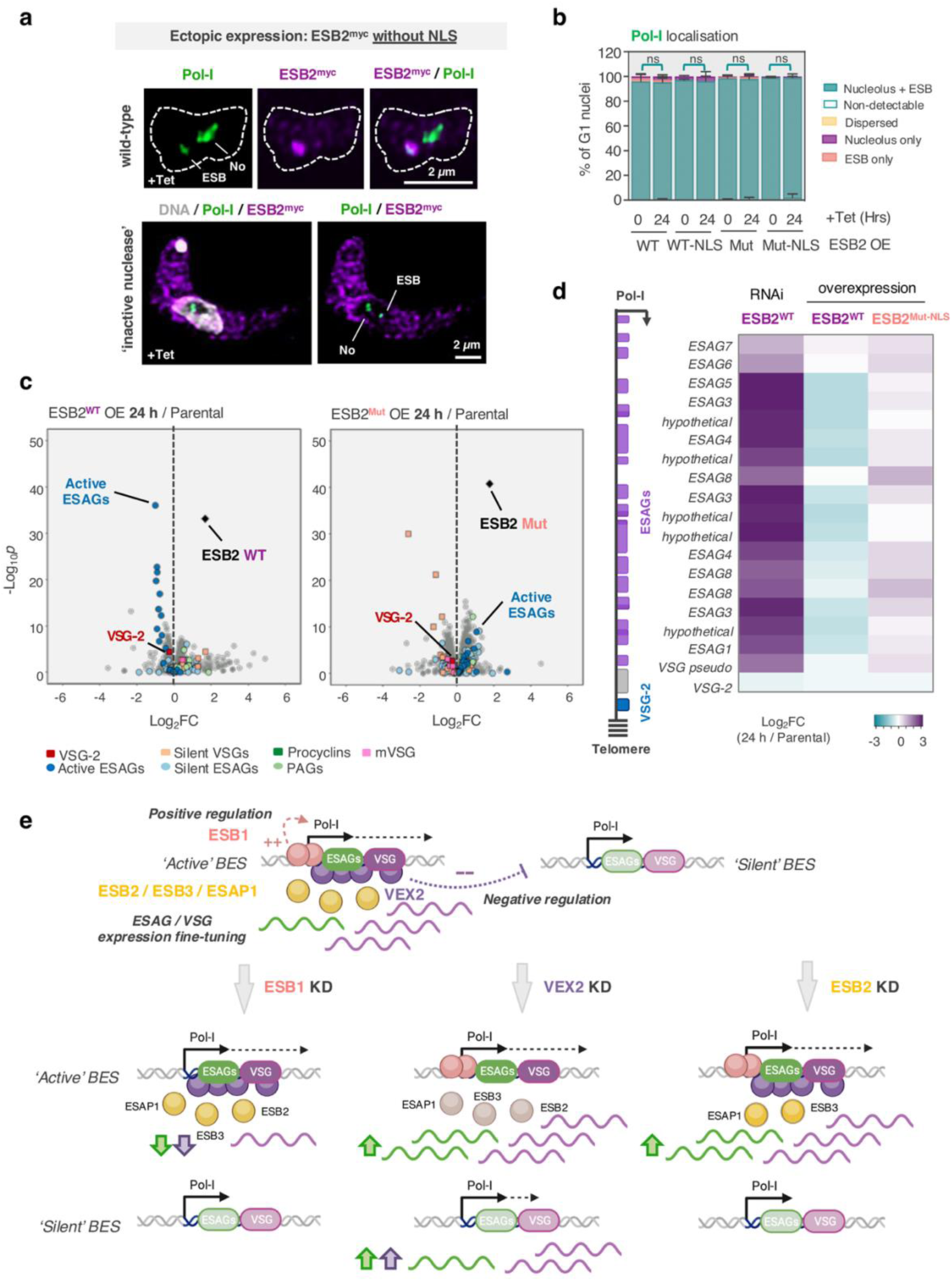
ESB2 fine-tunes *ESAG / VSG* expression at the ESB in a nuclease dependent manner. **a-b**, fluorescence microscopy analysis of Pol-I and ESB2^12myc^, wild-type or containing 3-point mutations (D240A, D330A, D353A) that render the nuclease inactive, with or without an La-NLS sequence. Images were acquired using a Zeiss LSM980 Airyscan 2 and correspond to 3D projections by brightest intensity of 0.1 μm stacks. DNA was stained with DAPI (grey); scale bars: 2 μm. The stacked graph (**d**) depicts averages of two biological replicates; >100 G1 cells per condition were analysed; error bars indicate standard deviation. Two-way ANOVA; ns, non-significant. The statistics depicted on the graph are focused on % of cells with an ESB (cyan). **c-d**, RNA-Seq analysis of ESB2^WT^ and ESB2^Mut-NLS^ overexpression (OE) at 72 hours post induction. Three biological replicates were used as well as parental and uninduced controls. **c**, Volcano plots depicting Log_2_FC (fold change) between overexpression and parental line and corresponding statistical significance (-log_10_*p*). The following cohorts are highlighted: ‘active’ *VSG* (red) and *ESAGs* (darker blue); ‘silent’ *VSGs* (orange) and *ESAGs* (lighter blue), metacyclic *VSGs* (mVSGs, pink), procyclins (darker green), procyclin associated genes (PAGs, lighter green). **d**, heatmap depicting the Log_2_FC within the active-ES (BES1) for induced samples of ESB2^WT^ and ESB2^Mut-NLS^ overexpression compared to ESB2 RNAi – all normalised against the parental line. From top to bottom, genes are ordered as they appear in BES1 (active-ES). *ESAG2* and *ESAG11* were omitted as their expression could not be detected in all replicates in the parental line when applying high stringency mapping. **e,** summary of the gene expression profile at both active and silent-ESs following ESB1, VEX2 or ESAP1/ESB3/ESB2 knockdowns. ESB1 is a transcriptional activator required for transcription at the active-ES. VEX2 is an exclusion factor, which prevents activation of silent *VSGs* and therefore a negative regulator. ESAP1 and ESB3 are required for ESB2 recruitment, which ultimately negatively regulates *ESAGs* expression. In the schematics, ESB2/ESB3/ESAP1 are not coloured in the VEX2 knockdown because they are compartmentalised in a VEX2-dependent manner. The upregulation of *ESAG* transcripts in the VEX2 RNAi is likely to be a consequence of ESB2 mislocalisation. Created with BioRender.com

